# Spatially clustered loci with multiple enhancers are frequent targets of HIV-1

**DOI:** 10.1101/287896

**Authors:** Bojana Lucic, Heng-Chang Chen, Maja Kuzman, Eduard Zorita, Julia Wegner, Vera Minneker, Vassilis Roukos, Wei Wang, Raffaele Fronza, Manfred Schmidt, Monsef Benkirane, Ralph Stadhouders, Kristian Vlahovicek, Guillaume J Filion, Marina Lusic

## Abstract

HIV-1 recurrently targets active genes that are positioned in the outer shell of the nucleus and integrates in the proximity of the nuclear pore compartment. However, the genomic features of these genes and the relevance of their transcriptional activity for HIV-1 integration have so far remained unclear. Here we show that recurrently targeted genes are delineated with super-enhancer genomic elements and that they cluster in specific spatial compartments of the T cell nucleus. We further show that these gene clusters acquire their location at the nuclear periphery during the activation of T cells. The clustering of these genes along with their transcriptional activity are the major determinants of HIV-1 integration in T cells. Our results show for the first time the relevance of the spatial compartmentalization of the genome for HIV-1 integration, thus further strengthening the role of nuclear architecture in viral infection.

## INTRODUCTION

Integration of the proviral genome into the host chromosomal DNA is one of the defining features of retroviral replication^1–3^. Upon infection of the target cell, the viral genome with viral and cellular proteins enters the nucleus as a pre-integration complex. As HIV-1 does not require cell division to enter the nucleus, the viral DNA passes through the nuclear pore complex (NPC) to access chromatin^4^. Nuclear pore proteins are important factors for the viral nuclear entry^5^, as well as for the positioning and consequent integration of the viral DNA into the cellular genome^3, 6^. In the T cell nucleus, HIV-1 resides predominantly in the outer spatial shells, associated with the open chromatin domains underneath the inner basket of the NPC^6^. However, the specific organization and features of the chromatin environment targeted by the viral DNA underneath the NPC remain unknown.

It is well established that HIV-1 integrates into active genes and regions bearing enhancer marks^7–9^. Unlike typical enhancers, genomic elements known as super-enhancers or “stretched enhancers” are defined by very high levels of activator binding or chromatin modifications, especially H3K27ac^10–12^. These clustered cis-regulatory elements are targeted by transcription factors different from the ones binding to typical enhancers, and include BRD4, the mediator complex^10^ and the p300 histone acetyltransferase^10, 12, 13^. Super-enhancers control the expression of genes that define cell identity, including many disease genes, tissue-specific genes or developmentally regulated genes^10, 12–14^. In the case of CD4^+^ T cells relevant for HIV-1 infection, super-enhancers were shown to control cytokines, cytokine receptors and transcription factors that regulate T cell specific transcriptional programs^15^. Strikingly, one of the strongest super-enhancers is associated with a gene encoding the transcription factor BACH2, a broad regulator of immune activation^16, 17^ and one of the most frequently targeted HIV-1 integration genes^18, 19^.

An additional feature of super-enhancer genomic elements and the genes that they control is the specific connection with nuclear pore proteins^20, 21^. In particular, Nup93 and Nup153 are involved in regulating the expression of cellular identity genes^20, 22^. Moreover, the binding of nucleoporins to super-enhancer elements is responsible for anchoring these genes to the nuclear periphery^21^. Super-enhancers seem to play a general role in organizing the genome through higher-order chromatin structures and architectural chromatin loops^23–25^. A disruption of higher-order chromatin structures, caused by depletion of the major architectural protein cohesin, results in disordered transcription of super-enhanced genes^23^. Super-enhancers, by contacting one another even when lying on different chromosomes, could anchor cohesin-independent architectural loops. This contributes to a new level of spatial compartmentalization and to the formation of so-called super-enhancer sub-compartment^23^.

Evidence accumulated in the last decade has revealed that the chromosomal contacts, achieved by genome folding and looping, define separate compartments in the nucleus^26^. Hi-C data have shown that transcribed genes make preferential contacts with other transcribed genes, forming a spatial cluster known as the A compartment^27, 28^. Reciprocally, silent genes and intergenic regions form a spatial cluster known as the B compartment. The loci of the B compartment are usually in contact with the nuclear lamina^29^, *i.e.* at the periphery of the nucleus. However, the active genes in contact with the nuclear pores are also peripheral, making nuclear periphery a composite environment, with features of either silent or active chromatin. Given the proposed location of super-enhanced genes in the NPC compartment^21, 23^ we were interested in understanding the relationships between HIV-1 integration sites, super-enhancers and genome organization.

We find that HIV-1 integrates in proximity of super-enhancers in patients and *in vitro* T cell cultures. The observed phenomenon does not depend on the activity of super-enhancers, but on their position in spatial neighborhoods where HIV-1 insertion is facilitated. Consistently, HIV-1 integration hotspots cluster in the nuclear space and tend to contact super-enhancers. Finally, we find that super-enhancer activity is critical to reorganize the genome of activated T cells, showing that they indirectly contribute to HIV-1 insertion biases.

## RESULTS

### HIV-1 integration hotspots are within genes delineated with super-enhancers

We assembled a list of 4,993 HIV-1 integration sites from primary CD4^+^ T cells infected *in vitro* (^30^ and this study) and 9,519 insertion sites from six published studies from HIV-1 patients^18, 19, 31–34^ (**Additional file 1**). 11,613 integrations were in gene bodies (85.5% for patient studies and 86% for *in vitro* infection studies), targeting a total of 5,824 different genes (**Supplementary Figure 1A**). This insertion data set is not saturating (**Supplementary Figure 1B**), but we found that a subset of genes are recurrent HIV-1 targets, consistent with our previous findings^6^. We thus defined Recurrent Integration Genes (RIGs) as genes with HIV-1 insertions in at least two out of eight datasets, yielding a total of 1,831 RIGs (**Supplementary Figure 1C**).

To characterize RIGs, we extracted genes without HIV-1 insertions in any dataset (called non-RIGs in the analysis; 51,939 genes in total) and compared their ChIP-Seq features in primary CD4^+^ T cells (**Figure 1A**). We observed higher levels of H3K27ac, H3K4me1 and H3K4me3, as well as BRD4 and MED1 at transcription start sites of RIGs vs non-RIGs. Histone profiles of H3K36me3 and H4K20me1 were higher throughout gene bodies of RIGs, while the repressive transcription mark H3K27me3 was lower on the gene bodies of RIGs vs non-RIGs. Of note, the mark of facultative heterochromatin H3K9me2 was depleted at transcription start sites of RIGs but remained unchanged throughout the gene body of RIGs vs non-RIGs.

**Figure 1.**
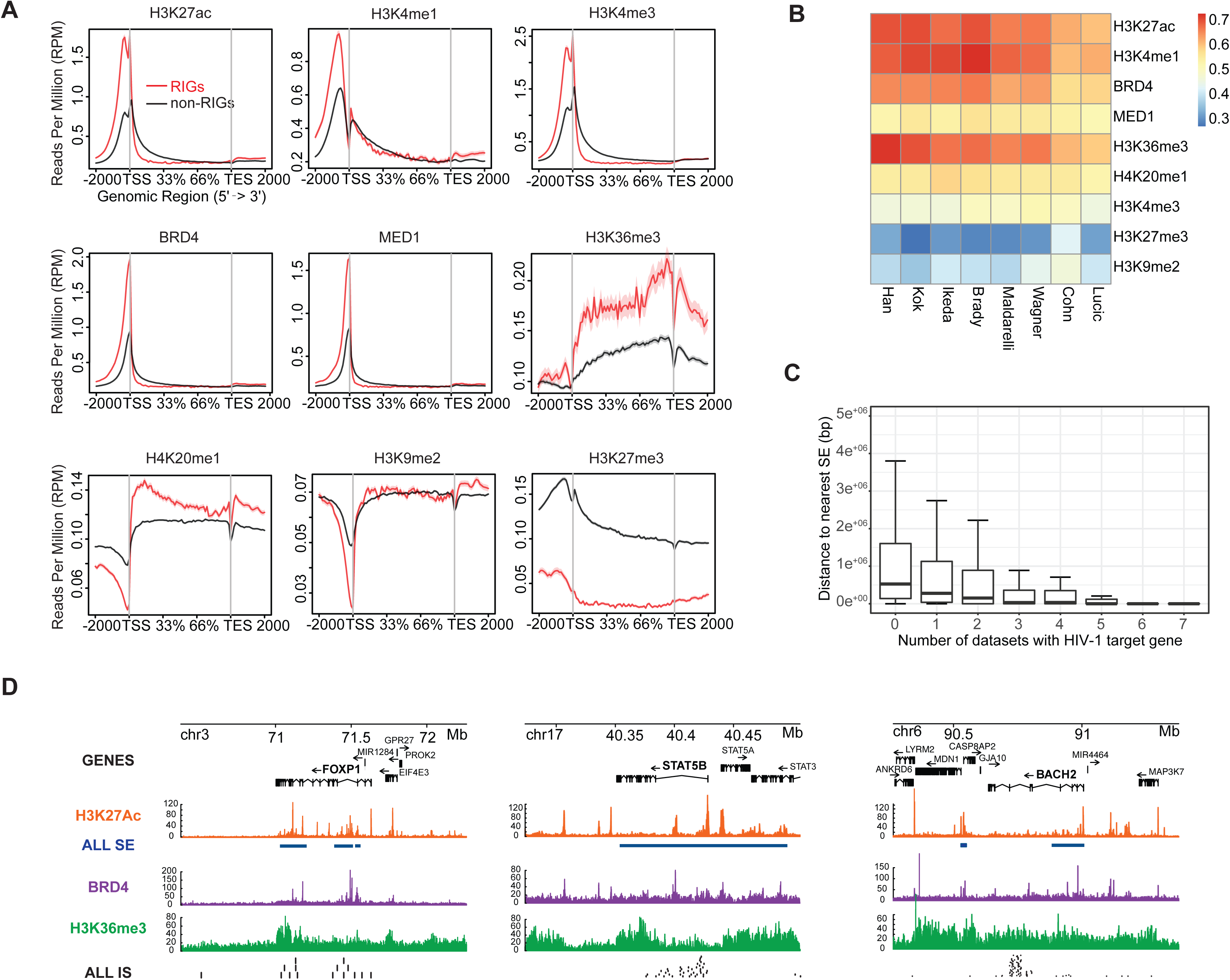
HIV-1 integration hotspots are within genes delineated with super-enhancers. **A)** Metagene plots of H3K27ac, H3K4me1, H3K4me3, BRD4, MED1, H3K36me3, H4K20me1, H3K9me2 and H3K27me3 ChIP-Seq signals in recurrent integration genes (RIGs) in red and the rest of the genes that are not targeted by HIV-1 (no RIGs) in black. **B)** ROC analysis represented in heatmap summarizing the co-occurrence of integration sites and epigenetic modification obtained by ChIP-Seq for H3K27ac, H3K4me1, BRD4, MED1, H3K36me3, H4K20me1, H3K4me3, H3K27me3 and H3K9me2. HIV-1 integration data sets are shown in the columns, and epigenetic modifications are shown in rows. Associations are quantified using the ROC area method; values of ROC areas are shown in the color key at the right. **C)** Distance to the nearest super-enhancer in activated CD4^+^ T cells. Box plots represent distance from the gene to nearest super-enhancer (SE) grouped by number of times gene is found in different data sets. **D)** *FOXP1*, *STAT5B* and *BACH2* ALL IS (black) superimposition on H3K27Ac (orange), SE (blue), H3K36me3 (green) and BRD4 (violet) ChIP-Seq tracks.

In order to test the specificity of chromatin signatures of HIV-1 integration sites, we adapted the receiver operating characteristic (ROC) analysis^35, 36^. We used controls sites matched according to the distance to the nearest gene (see Methods) and confirmed significant enrichment reproducible over all datasets of the following genomic features: H3K27ac, H3K4me1, BRD4, MED1, H3K36me3 and H4K20me1 (**Figure 1B**). The marks H3K27ac, H3K4me1 and H3K36me3, characteristic of active enhancers^37^, cell type specific enhancers^38^, and bodies of transcribed genes^39^, respectively, were the most enriched in the proximity of insertion sites. Consistent with the presence of H3K27ac and H3K4me1, we also found significant enrichment of BRD4, a constituent of super-enhancer genomic elements^10, 12^ (**Figure 1B**). On average, 60% of insertion sites were significantly enriched in these chromatin marks (not shown) while we observed depletion of H3K27me3 and H3K9me2 in the proximity of insertion sites. Interestingly, we did not observe a statistically significant enrichment of H3K4me3 in the proximity of insertion sites.

We therefore identified super-enhancers in activated CD4^+^ T cells using H3K27ac ChIP-Seq and merged them with the super-enhancers in activated CD4^+^ T cells from dbSuper^40, 41^. We obtained 2,584 super-enhancers, intersecting 360 RIGs out of 1,813 (34.41%, **Supplementary Figure 1D**). In addition, the more a RIG is targeted by HIV-1 (*i.e.* the higher the number of datasets where HIV-1 insertions are found in the gene), the closer it lies to super-enhancers on average (**Figure 1C**). In contrast, the insertion sites of the retrovirus HTLV-1^42^ (Human T Lymphotropic Virus type 1) were not enriched in super-enhancer marks (**Supplementary Figure 1E**). **Figure 1D** shows the integration biases at gene scale on *FOXP1*, *STAT5B* and *BACH2*, three highly targeted RIGs involved in T cell differentiation and activity. The ChIP-Seq profiles of H3K27ac, H3K36me3 and BRD4 indicate prominent clustering of HIV-1 insertion sites near the super-enhancers defined by those marks. Thus, HIV-1 tends to integrate into genes delineated with super-enhancers, a tropism that is not a general feature of retroviruses.

### RIGs are delineated with super-enhancers regardless of their expression

HIV-1 is known to integrate into highly expressed genes^7, 9^. It is thus possible that genes with a super-enhancer are targeted more often because they are expressed at higher level. To test whether this is the case, we measured the transcript abundance of genes in CD3/CD28-activated CD4^+^ T cells by RNA-Seq. The mean expression of genes with HIV-1 insertions is higher than those not targeted by HIV-1 (**Figure 2A**). More specifically, 74.8% of genes targeted by HIV-1 are in the “high expression” group, compared to 52.8% of non-targeted genes (see Methods). Moreover, the genes more often targeted by HIV-1 (RIGs) are expressed at higher levels (**Figure 2B**), thus confirming that HV-1 is biased towards highly expressed genes.

**Figure 2.**
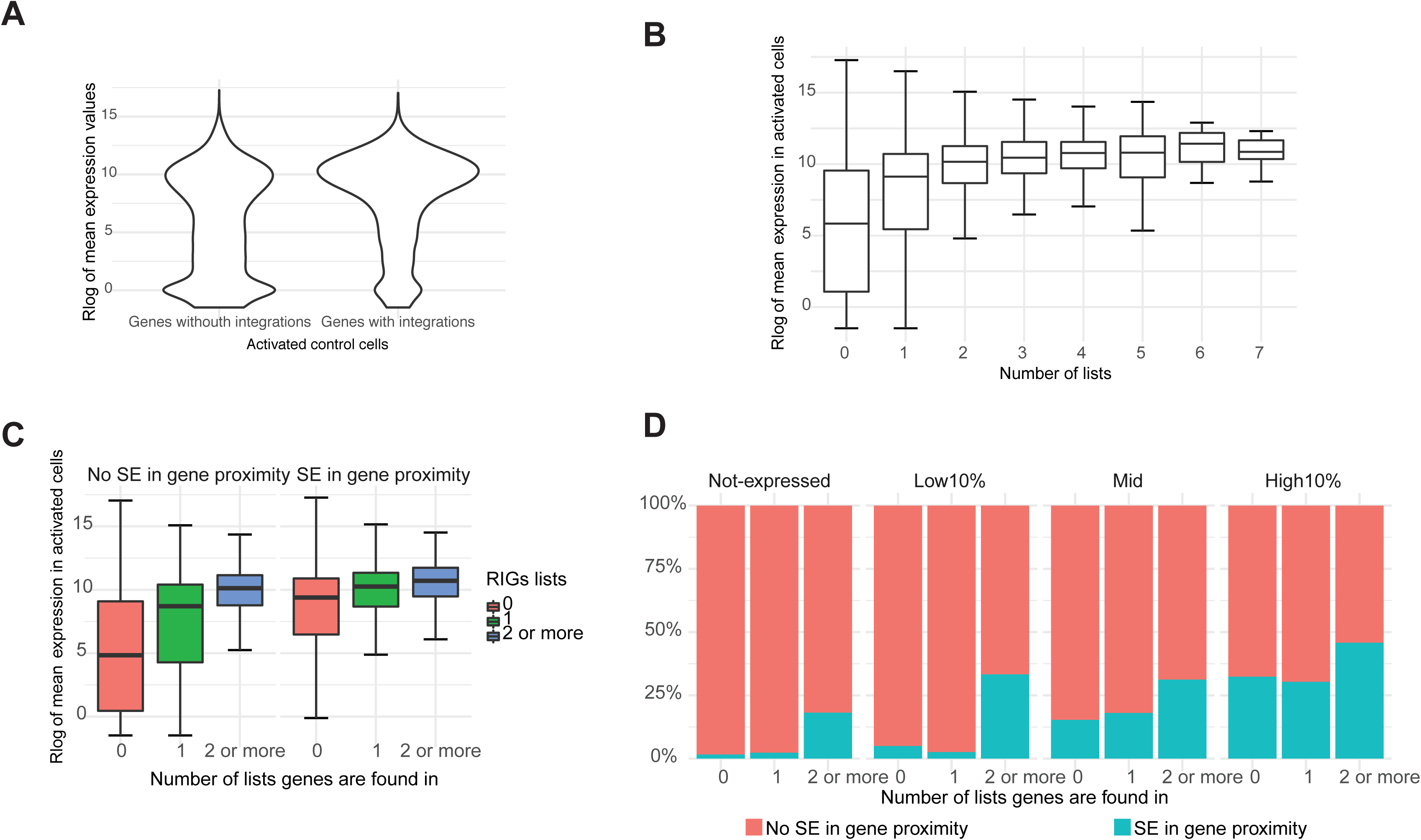
RIGs are delineated with super-enhancers regardless of their expression. **A)** Regularized log transformed read counts on genes averaged over three replicates in activated CD4^+^ T cells shown as violin plot for genes without HIV-1 integrations and genes with HIV-1 integrations. **B)** Boxplot for genes grouped by number of HIV-1 lists they appear in. **C)** Boxplot for genes grouped by number of HIV-1 lists they appear in, with RIGs grouped together in 2 or more lists group. Boxplots are shown separately for genes that have super-enhancer 5 kb upstream of TSS or super-enhancer overlaps them (SE in proximity), and genes that do not have super-enhancer in proximity. Differences in median abundances of mRNA are statistically significant for all groups (p-value < 2.2×10^-16^ for genes without HIV integrations and genes found on only one list and p-value 3.7×10^-12^ for RIGs, calculated by Wilcoxon rank sum test). **D)** Bar plot shows percentage of genes that have super-enhancer in proximity, divided by number of lists gene is found in and expression group.

On average, genes with a super-enhancer are expressed at higher levels than those without (**Figure 2C**). This trend is more subtle for RIGs, as they are expressed at high level, with or without super-enhancers (**Figure 2C**, compare the blue boxes). However, RIGs are more often in the proximity of super-enhancers than non-RIGs, irrespective of their expression (**Figure 2D**). In particular, 18.2% of RIGs that are silent also have a proximal super-enhancer, while this is true for only 1.7% of the silent genes that were never found to be HIV-1 targets (**Figure 2D**, leftmost panel). The trend remains the same for expressed genes (**Figure 2D**) after dividing them in “low”, “medium”, and “high” expression groups (see Methods).

### Super-enhancer unloading in activated CD4^+^ T cells does not impact HIV-1 integration patterns

To assess the relationship between HIV-1 integration and transcription of genes controlled by super-enhancer elements, we took advantage of the BET inhibitor JQ1, which unloads BRD4 from chromatin^43^ and causes a subsequent dysregulation of RNA Pol II binding^11^.

We compared the HIV-1 insertion profiles with or without JQ1 in CD4^+^ T cells, where RNA and protein levels of *MYC* (a bona fide super-enhancer gene^11^) served as a control (**Supplementary Figure 2A**). Using inverse PCR (see Methods), we mapped a total of 38,964 HIV-1 insertion sites. At the chromosome scale, the insertion biases are not affected by JQ1 (**Figure 3A**, left panel). In particular, the characteristic ∼3-fold enrichment on chromosomes 17 and chromosome 19 was not perturbed. The right panel of **Figure 3A** represents an “insertion cloud” on chromosome 17, where each mapped HIV-1 insertion is plotted at its position on the x-axis, but at a random position on the y-axis. This way, insertion hotspots appear as vertical lines, domains enriched in insertions appear as bursts, and insertion deserts appear as empty spaces. For comparison purposes, the insertion profile upon JQ1 treatment was flipped vertically. The profiles with or without JQ1 are similar, and the hotspots are not perturbed by the treatment, indicating that the insertion biases were maintained upon treatment. In addition, we did not observe any statistically significant change of insertion pattern in genes vs intergenic regions, nor in other types of genomic regions (data not shown).

**Figure 3.**
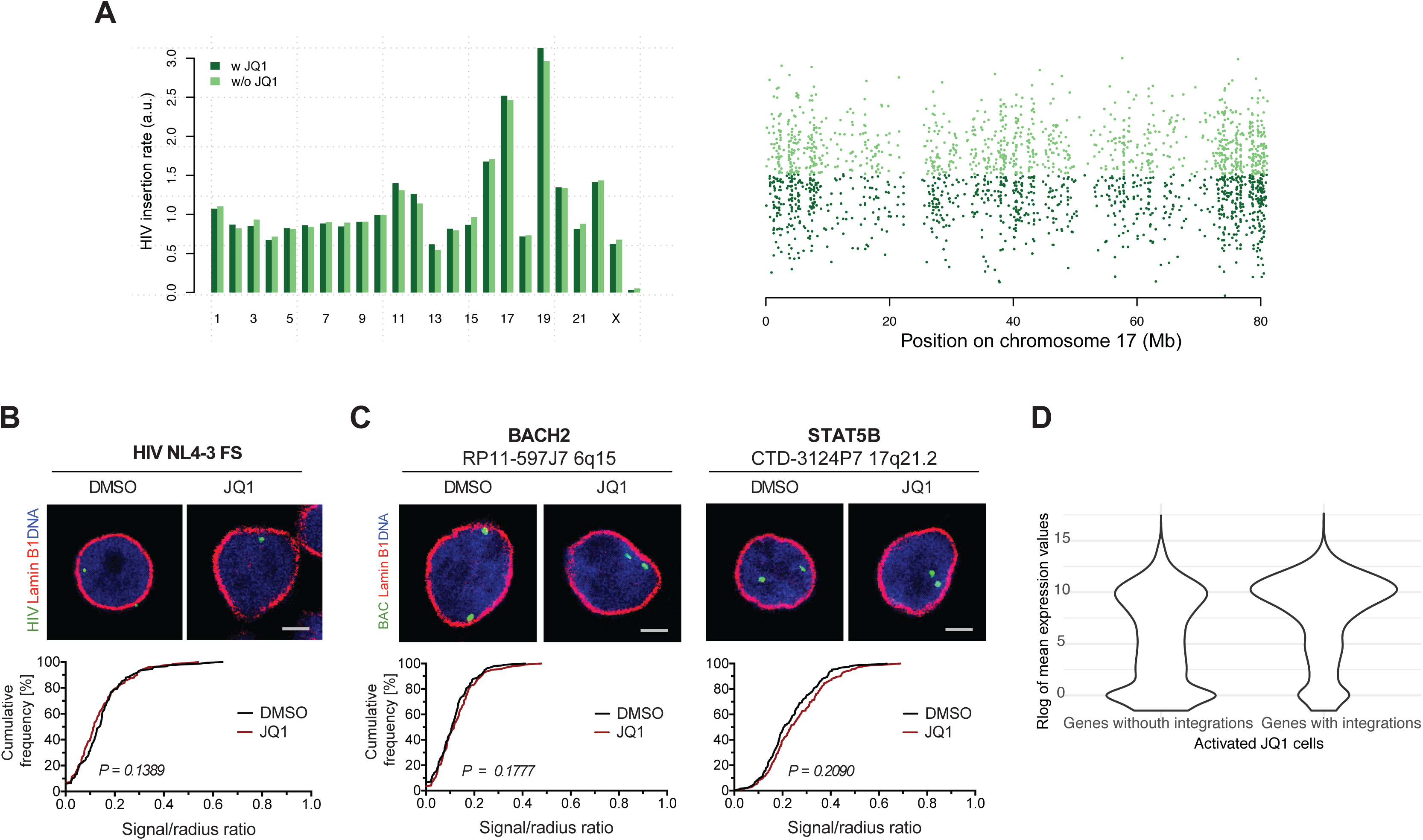
Super-enhancer unloading in activated CD4^+^ T cells does not impact HIV-1 integration patterns. **A)** Bar plot of HIV-1 insertion rate per chromosome: control (w/o JQ1) or JQ1 treated (w JQ1) samples (left panel) and ‘insertion cloud’ representation on chromosome 17 (right panel). Each dot represents an HIV-1 insertion site. The x-coordinate indicates to the location of the insertion site on chromosome 17; the y-coordinate is random so that insertion hotspots appear as vertical lines. **B)** 3D immuno-DNA FISH images of HIV-1 in activated CD4^+^ T cells pretreated with 500 nM JQ1 for 6 h and infected for 72 h (green: HIV-1 probe, red: lamin B1, blue: DNA staining with Hoechst 33342, scale bar represents 2 μm). Cumulative frequency plots show combined data from both experiments (n = 100, black: DMSO, red: JQ1). The p-values of the Kolmogorov-Smirnov tests are indicated. **C)** 3D immuno-DNA FISH images of *BACH2* and *STAT5B* upon 500 nM JQ1 treatment for 6 h in activated CD4^+^ T cells (green: BAC/gene probe, red: lamin B1, blue: DNA staining with Hoechst 33342, scale bar represents 2 μm). **D)** Regularized log transformed read counts on genes averaged over three replicates in activated JQ1 treated cells shown as violin plot for genes grouped by presence of HIV-1 integration in activated JQ1 treated cells.

We used 3D FISH to test whether the localization of the provirus in the nuclear space was affected, but did not observe any significant difference (**Figure 3B**). Consistently, the positions of two RIGs with super-enhancers, *BACH2* and *STAT5B*, were also not affected by the JQ1 treatment (**Figure 3C**). Next, we profiled the transcriptome of activated CD4^+^ T cells and observed, independently of HIV-1 integration, that more genes proximal to super-enhancers show significant up- or down-regulation upon JQ1 treatment than genes without a super-enhancer (**Supplementary figure 2B** and **2C**). When we specifically looked at genes targeted by HIV-1, we observed that the effect of JQ1 is even more pronounced among RIGs than among not-targeted genes (**Supplementary figure 2C**). By comparing the integration and transcription profiles we observe that the global insertion bias towards active genes is not modified by the JQ1 treatment (compare **Figure 2A** and **Figure 3D**). Hence, HIV-1 retains the patterns of integration into highly transcribed genes in both control and JQ1 treated cells.

### HIV-1 integration hotspots are clustered in the nuclear space

Our previously published results showed that RIGs are distributed in the periphery of the T cell nucleus^6^, so we hypothesized that the enrichment of HIV-1 insertion sites near super-enhancers may be due to their particular organization in the nuclear space. We thus performed Hi-C to get some insight into the conformation of the T cell genome.

In order to minimize reproducibility issues caused by the heterogeneity of the biological material, we used the widely available Jurkat lymphoid T cellular model. To ensure that the behavior of HIV-1 is similar in both models, we compared a published collection of 58,240 insertion sites in Jurkat cells^8^ to the 28,419 insertion sites in primary CD4^+^ T cells form this and previous studies^30^. The insertion rates per chromosome are similar between cells (**Figure 4A**); both show the characteristic ∼3-fold increase on chromosomes 17 and 19. It is possible that the apparent difference on chromosome 17 is due the use of different mapping technologies. For comparison, our previous measure of the insertion rates on chromosome 17 of Jurkat cells^9^ is very close to the current measure in primary CD4^+^ T cells (*i.e.* using the same inverse PCR technology). The insertion cloud representation shows that the profiles are similar on chromosome 17, with the exception of a hot spot visible only in primary CD4^+^ T cells at ∼57 Mb (**Figure 4B**). We also found that the HIV-1 target genes are similar in Jurkat cells and in other CD4^+^ data sets (**Supplementary Figure 3**). In summary, apart from minor differences, HIV-1 insertion biases are comparable in primary CD4^+^ T and in Jurkat cells.

**Figure 4.**
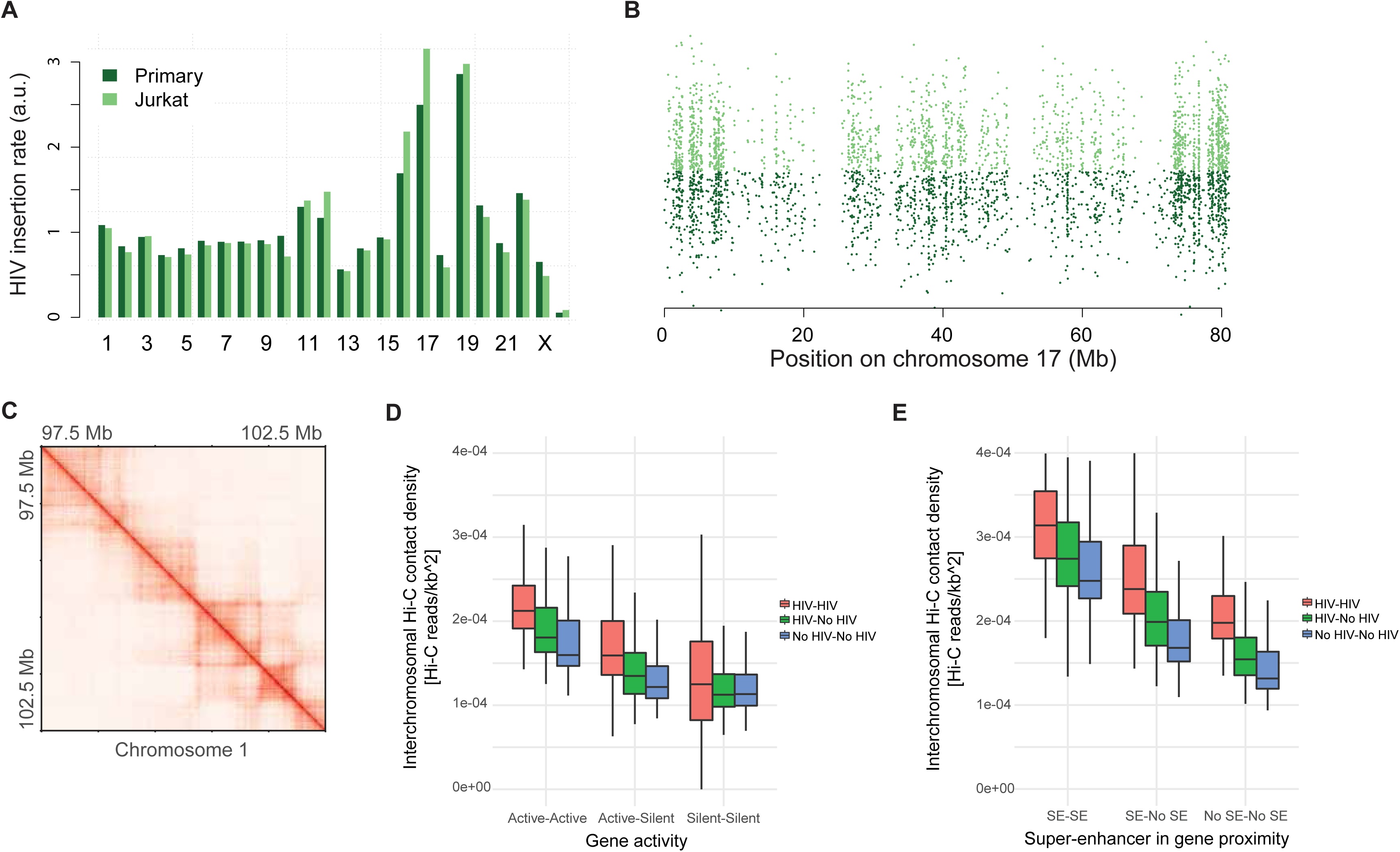
HIV-1 integration hotspots are clustered in the nuclear space. **A)** Bar plot of HIV-1 insertion rate per chromosome (the genome-wide average is set to 1) in primary T and in Jurkat cells. **B)** HIV-1 insertion cloud on chromosome 17 in primary T and Jurkat cells. Each dot represents an HIV-1 insertion site. The x-coordinate indicates to the location of the insertion site on chromosome 17; the y-coordinate is random so that insertion hotspots appear as vertical lines. **C)** Detail of the Hi-C contact map in Jurkat in 5 kb bins. TADs and loop domains are clearly visible. **D)** Box plot of inter-chromosomal Hi-C contact density (see Methods). Contact densities were computed between chromosomal aggregates of all gene fragments (5 kbp) corresponding to Active and Silent genes, with (HIV) or without HIV insertions (No HIV). The distribution of densities are composed of the scores for all inter-chromosomal combinations. **E)** Same as in D, but genes are classified between genes in proximity of super enhancers (SE), i.e. within gene body or 5 kbp upstream of TSS, or far from super-enhancers (No SE).

Hi-C on uninfected Jurkat cells yielded ∼1.5 billion informative contacts. Topologically Associating Domains (TADs) and loop domains are clearly visible on the raw Hi-C map in 5 kb bins (**Figure 4C**), showing that the experiment captures the basic structural features of the Jurkat genome. We also verified that the A and B compartments are well defined and that they correspond to the regions of high and low gene expression, respectively (data not shown). To our knowledge, this dataset constitutes the highest resolution Hi-C experiment presently available in Jurkat cells. If the insertion pattern of HIV-1 reflects a particular organization of the genome, one predicts that the insertion hotspots occupy the same nuclear space and thus cluster together in 3D. We tested this hypothesis by measuring the amount of inter-chromosomal Hi-C contact densities among different classes of HIV-1 insertion sites (**Figure 4D**). The loci most targeted by HIV-1 engage in stronger contact with each other than non-targeted loci. Also, the differences in contact strength are more pronounced when such genes are active. In addition, super-enhancers tend to cluster together and with HIV-1 insertion hotspots in 3D (**Figure 4E**, **Supplementary Figure 4**), indicating that super-enhancers locate in the physical proximity of HIV-1 insertion sites. Thus, HIV-1 insertion sites form spatial clusters interacting with super-enhancers in the nucleus, consistently with the view that the insertion process depends on the underlying 3D organization of the T cell genome.

### Super-enhancers and HIV-1 occupy the same 3D sub-compartment

To better define the properties of HIV-1 insertion sites, we segmented the Jurkat genome into spatial clusters. For each chromosome, we generated 15 clusters of loci enriched in self interactions, that we coalesced down to 5 genome-wide clusters based on their inter-chromosomal contacts (**Figure 5A** and see Methods). This approach yielded two A-type sub-compartments called A1 and A2, two B-type sub-compartments called B1 and B2, and one intermediate / mixed compartment called AB (**Figures 5B and 5C**).

**Figure 5.**
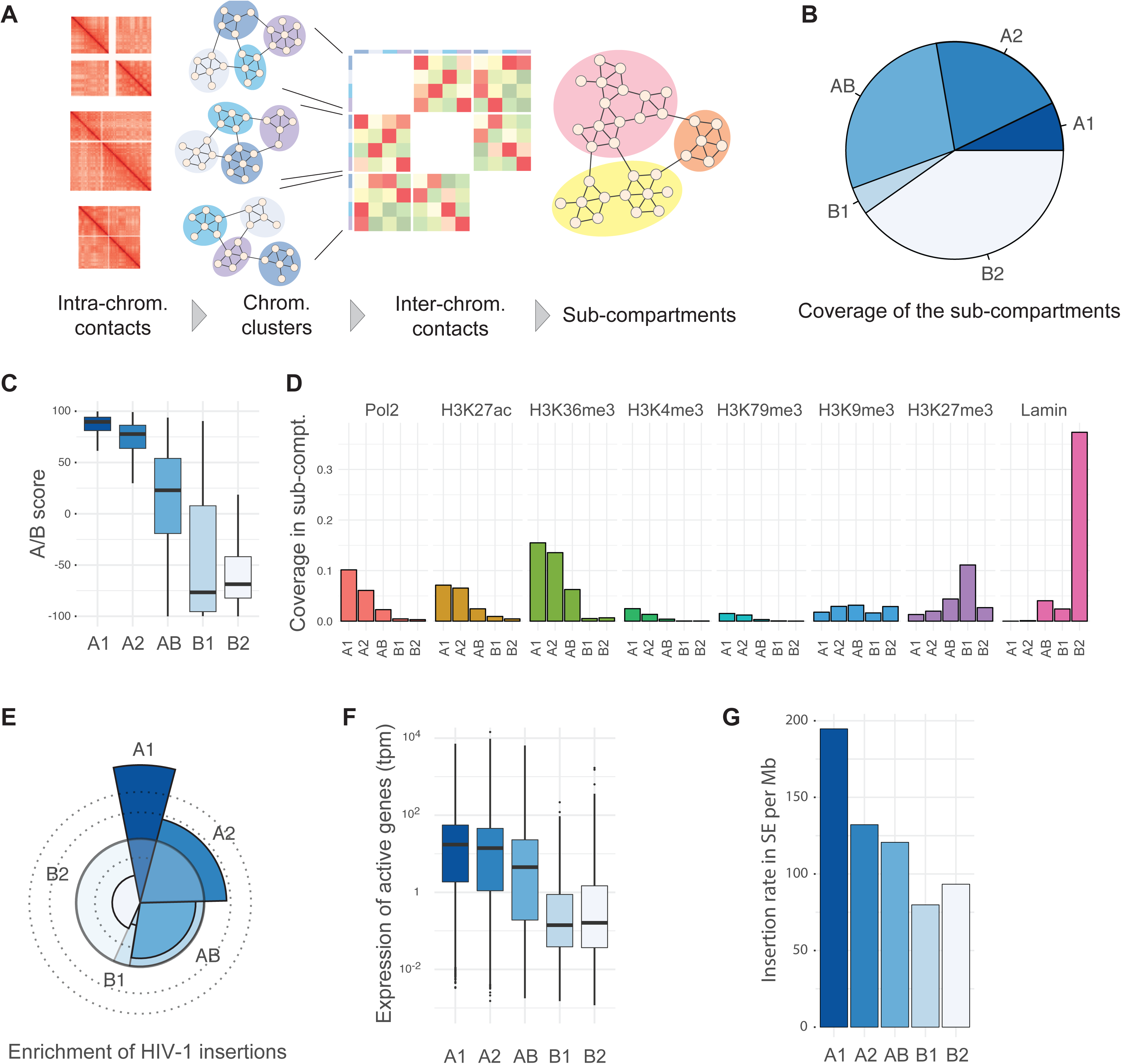
Super-enhancers and HIV-1 occupy the same 3D sub-compartment. **A)** Definition and identification of 3D compartments. For each chromosome, 15 spatial communities were identified by clustering. The inter-chromosomal contacts between the communities were used as a basis for another round of clustering in 5 genome-wide spatial communities. **B)** Pie chart showing the coverage of the sub-compartments in the Jurkat genome. **C)** Distribution of AB scores in the 3D sub-compartments. The AB score measures the likelihood that a locus belongs to the A or B compartment. Extreme values +100 and −100 stands for “fully in A” or “fully in B”, respectively. A score of 0 means “both or neither”. **D)** Proportion of 3D sub-compartments covered by major chromatin features. Coverage was computed as the span of enriched ChIP-Seq signal divided by the sub-compartment size. **E)** Spie chart showing the observed vs expected HIV-1 insertions in the sub-compartments. The expected amount of insertions is the area of the wedge delimited by the circle in bold line, and the observed amount is the area of the colored wedge. Dotted lines represent the limit of the wedge for 2x and 3x enrichment outside the circle, and 0.5x depletion inside the circle. The observed / expected ratio is approximate 2.5 times higher in A1 than in A2. **F)** Box plot showing the expression of active genes in the sub-compartments. The plot was rendered using defaults from the ggplot2 library. **G)** Bar plot showing the integration density of HIV in super-enhancers located in different 3D sub-compartments.

The B-type sub-compartments correspond to known types of silent chromatin. B1 is enriched in the Polycomb mark H3K27me3, and B2 is enriched in Lamin (**Figure 5D** and **Supplementary Figure 5A**). The AB sub-compartment is marked by features of euchromatin and heterochromatin, each at low coverage. This suggests that the AB sub-compartment is composed of loci at the transition between euchromatin and heterochromatin. Finally, the A-type sub-compartments are both enriched in euchromatin marks, with higher coverage in A1 than in A2 (**Figure 5D** and **Supplementary Figure 5A**).

Strikingly, the rate of HIV-1 insertion is ∼2.5 times higher in A1 than in A2 (**Figure 5E**). In contrast, the coverage of euchromatin marks and the transcriptional activity are only slightly higher in A1 than in A2 (**Figures 5D and 5F**). More importantly, the ∼2.5-fold enrichment is still present when controlling for gene expression (**Supplementary Figure 5B**), indicating that HIV-1 insertion is intrinsically more frequent in the A1 sub-compartment. Of note, we obtained similar results when defining 10 sub-compartments instead of 5, where HIV-1 insertion rates are enriched in one sub-compartment covering ∼10% of the genome (data not shown). Hence, our observation is robust with respect to the definition of sub-compartments. The 3D organization of the Jurkat T cell genome thus explains large differences of HIV-1 insertion rates between genes expressed at similar levels.

If HIV-1 targets super-enhancers because of their location in the nuclear space, one predicts that the insertion rate of HIV-1 in the super-enhancers of A1 should be higher than in the super-enhancers of A2. **Figure 5G** shows that, indeed, HIV-1 is ∼1.5 times more likely to integrate in the super-enhancers of A1 than in those of A2. Since the insertion rate in super-enhancers depends primarily on their location, we conclude that the enrichment in super-enhancers at genome-wide scale is due to their position in the 3D space of the nucleus, rather then due to their activity or their chromatin features.

### Super-enhanced genes relocate to the nuclear periphery upon T cell activation

Our results so far suggest that HIV-1 hitchhikes on super-enhancers because of their location in the structured genome of T cells, but they do not address the contribution of super-enhancers to this structure. We thus investigated the role of super-enhancers in the transition from resting to activated T cells. For this, we used primary T cells.

RIGs belong to a subset of T cell genes that show the strongest response to T cell activation (**Supplementary Figure 6A**). As RIGs are rich in super-enhancers and as they are located at the nuclear periphery in activated CD4^+^ T cells^6^, we reasoned that their spatial positioning might change with the activation status of cell. We therefore employed 3D immuno-DNA FISH to visualize gene positioning in resting and activated CD4^+^ T cells. The cumulative frequency plots revealed that nine RIGs, seven of which have super-enhancers *FOXP1, STAT5B, NFATC3* (**Figure 6A),** *KDM2A, PACS1* (**Figure 6B)**, and *GRB2, RNF157* (**Figure 6C)**, shifted upon activation towards the periphery of the T cell nucleus. Two non-RIGs were used as controls: *PTPRD*, positioned at the periphery, and *TAP1*, positioned at the center of the nucleus of activated CD4^+^ T cells^6^. In resting cells, *PTPRD* was equally peripheral and *TAP1* was equally central (**Supplementary figure 6B**). Thus, in contrast to the tested RIGs, both controls did not change position upon CD4^+^ T cell activation.The observed gene movement appears to be a general feature of genes with super-enhancers, since also the non-RIG *MYC* harboring five well-described super-enhancers changed radial positioning (**Supplementary Figure 6C**). Of note, three RIGs, *NPLOC4* (found in 4 data sets) *RPTOR* (in 7 data sets) and *BACH2* (in 6 data sets) maintained their peripheral nuclear localization in both resting and activated states (**Supplementary figure 6E**).

**Figure 6.**
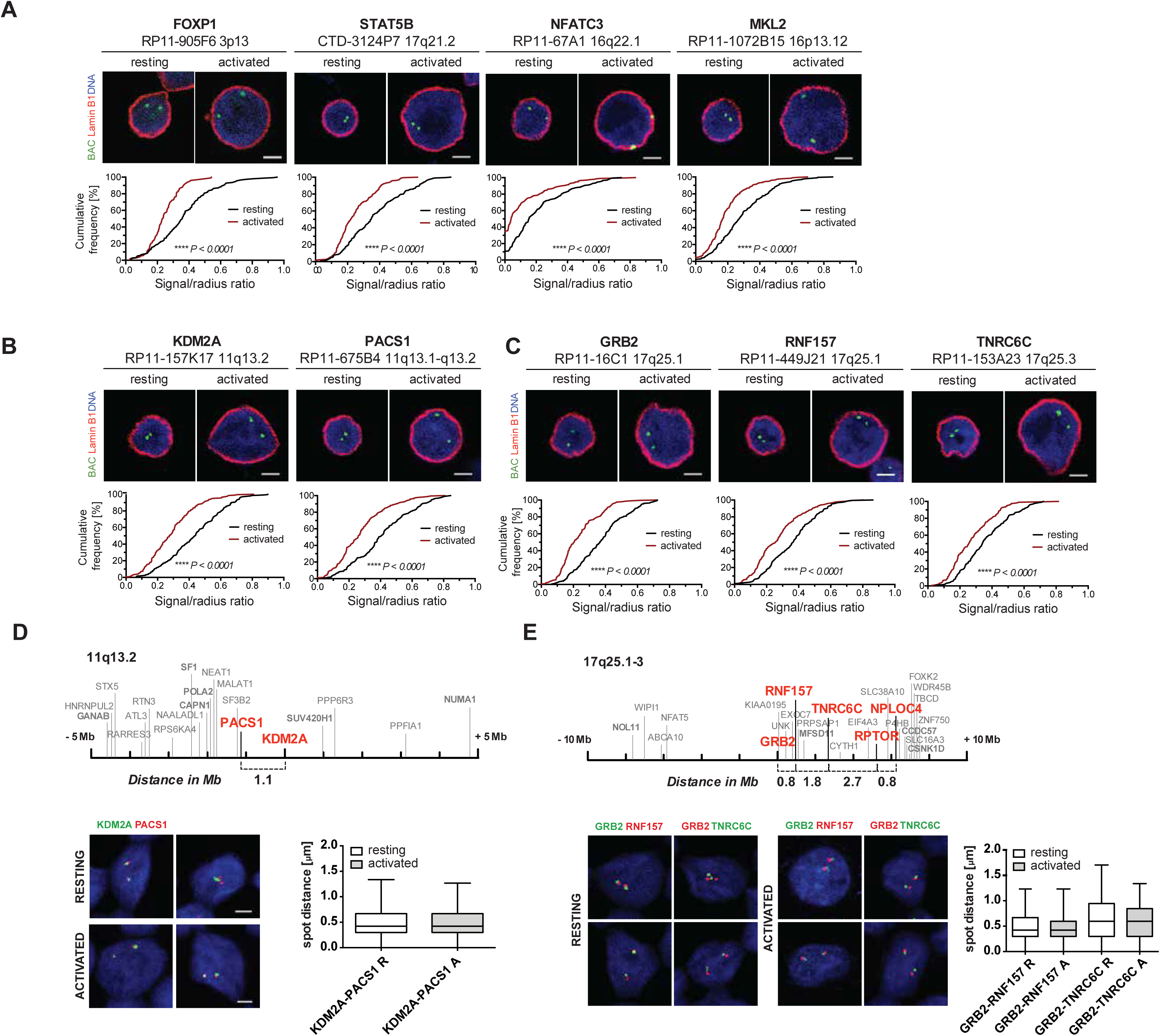
Super-enhancer associated genes relocate to the nuclear periphery upon T cell activation. Three-dimensional immuno-DNA FISH of nine RIGs in resting and activated (anti-CD3/anti-CD28 beads, IL-2 for 48 h) CD4^+^ T cells (green: BAC/gene probe, red: lamin B1, blue: DNA staining with Hoechst 33342, scale bar represents 2 μm). Cumulative frequency plots show combined data from both experiments (n = 100, black: resting cells, red: activated cells). The p-values of the Kolmogorov-Smirnov tests are indicated. Box plots represent minimized distances (5-95 percentile) for the analyzed gene combinations in resting (white) and activated (gray) CD4^+^ T cells, obtained by high throughput imaging and subsequent computational measurements. **A)** Representative images for *FOXP1*, *STAT5B*, *NFATC3* and *MKL2*. **B)** *KDM2A* and *PACS1*. **C)** *GRB2*, *RNF157* and *TNRC6*C. **D)** Schematic representation of chromosomal region 11q13.2 within 10 Mb: RIGs (bold red) and single HIV-1 integration sites (plain gray) and HTI of *KDM2A* and *PACS1*. **E)** Schematic representation of chromosomal region 17q25.1-3 within 20 Mb: RIGs (bold red) and single HIV-1 integration sites (plain gray) and HTI of *GRB2*, *RNF157* and *TNRC6C*.

We next selected two highly targeted regions on chromosomes 11 and 17 with multiple RIGs (percentage of RIGs among the top 10% of all regions of same size). By employing a dual-color FISH coupled to high-throughput imaging^44, 45^ we analyzed ∼10,000 alleles in 5000 cells in both resting and activated states. We observed that two genes from chromosome 11, *KDM2A* and *PACS1*, separated at the linear genomic scale by 1.1 Mb (**Figure 6D**), clustered together in the nuclear space, with a minimized median distance of 0.42 μm (data summarized in **Supplementary Figure 6E**).

Such spatial distribution is not always observed among genes (RIGs) that are close on a linear scale. When compared to other two RIGs, *NPLOC4* and *RPTOR* from Chromosome 17, clustering was not observed despite them being only 0.8Mb apart (**Supplementary Figure 6E,** images not shown). Interestingly, this region mapping to q25.1-3 on Chromosome 17 contains other 23 RIGs (**Figure 6E**). In contrast to *NPLOC4* and *RPTOR*, three RIGs, *GRB2*, *TNRC6C* and *RNF157*, cluster together (**Figure 6E**), as shown by using different combinations of FISH probe for dual-color FISH coupled to high-throughput imaging. Their spatial relationships did not change with the activation status of CD4^+^ T cells (**Figure 6E**, data summarized in **Supplementary Figure 6E**). These data suggest that RIGs and genes with super-enhancers relocate to the nuclear periphery in clusters (**Figures 6D and 6E**). As the radial positioning of genes with super-enhancer genomic elements changes with the activation status of CD4^+^ T cells, we wanted to understand if loading of super-enhancers impacts the nuclear position of super-enhanced genes. We assessed the 3D nuclear position of two RIGs with super-enhancer elements in resting T cells and compared them with T cells treated with JQ1 prior to the cellular activation with CD3/CD28 antibodies. Although the activation status of cells pretreated with JQ1 does not differ from the normally activated T cells (data not shown), we observed that two of the super-enhanced RIGs retain their position in the center of the nucleus, *i.e.* do not shift towards the periphery (**Supplementary Figure 6F**).

As we find that seven out of nine RIGs delineated with super-enhancers change radial positioning in activated CD4^+^ T cells we conclude that radial positioning of genes with super-enhancers changes with the activation status of CD4^+^ T cells. Overall, our results indicate that during T cell activation, super-enhancers poise the genome into a particular structure that defines the insertion biases of HIV-1. Genes with a super-enhancer move to the periphery of the nucleus, where they are exposed to higher insertion rate upon HIV-1 infection.

## DISCUSSION

The integration of the viral DNA into the host cell genome is responsible for the long term persistence of HIV-1 in cellular reservoirs^46^. The persistence of HIV-1 is influenced by the chromosomal context at the sites of integration, with a strong impact on the outcome of viral infection^9^. Here, we characterized the genomic features of integration sites identified from patients^18, 19, 31–34^ and from *in vitro* infections of CD4^+^ T cells (^30^ and this study). By analyzing these large datasets we confirmed that HIV-1 repeatedly integrates into a subset of active cellular genes, which we found to be delineated with super-enhancer genomic elements (**Figure 1C**). Yet, neither the activity of super-enhancers, nor their effect on gene expression alone explain the integration process (**Figure 3A**). Instead, we found that the correlation can be attributed to the enrichment of super-enhancers in the A2 and especially the A1 sub-compartments, where HIV-1 integrates at higher frequency than in the rest of the genome (**Figure 5E**). Super-enhancers do not affect the HIV-1 integration process once the genome structure is established, but we found that during T cell activation, they anchor their target genes at the periphery of the nucleus (**Figures 6A-C**).

The contribution of gene expression levels to the insertion rate of HIV-1 is intricate. On one hand, HIV-1 shows a clear bias towards expressed genes, even upon JQ1 treatment, where HIV-1 still integrates in active genes. However, HIV-1 recurrently integrates into genes with super-enhancers (**Figure 2D**), among which there are both up- and down-regulated genes (**Supplementary figure 2C**). One potential explanation is that the HIV-1 integrase has a strong affinity for some protein present in transcribed regions (*e.g.*, LEDGF). The complete absence of such proteins in non-transcribed regions would have more influence on the signal than its quantitative variations in transcribed regions. In any event, gene expression and chromatin are not the sole contributors to HIV-1 insertion patterns. The A1 sub-compartment is targeted more frequently than the rest of the genome (**Figure 5E**), even when controlling for chromatin and gene expression (**Supplementary Figure 5B**). This indicates that the 3D genome organization of activated T cells is an important determinant of the HIV-1 insertion process.

Does the sub-compartment A1 correspond to loci at the nuclear pore? It is indeed tempting to speculate that this is the case because we previously showed that these sites are frequent targets of HIV-1^6^. In IMR90 and U2OS cells, nuclear pore proteins are enriched in the A1 sub-compartment defined here (^21^ and data not shown) but it remains to be determined whether this is also the case in Jurkat or primary CD4^+^ T cells. None of the chromatin features mapped in Jurkat cells is known to discriminate active genes at the nuclear pore from other active genes, and the chromatin of A1 is otherwise similar to that of A2 (**Supplementary Figure 5A**). Interestingly, the density of super-enhancers is similar between A1 and A2 (**Supplementary Figure 5A**), so it is unlikely that the A1 sub-compartment simply emerges from the clustering of super-enhancers. More plausibly, super-enhancers are one of many contributors to the segregation of the genome in spatial clusters. More generally, the existence of two separate clusters of active genes in the 3D space of the nucleus is itself an intriguing observation that will require more work to be fully understood.

The clustering of the genome and higher-order chromatin structures, in particular architectural loops were recently linked to super-enhancers^23^. A transient complete loss of the main architectural protein cohesin leads to disruption of higher-order chromatin structures. At the same time, this results in the formation of higher-order hubs containing multiple super-enhancers and in the disordered transcription of genes predicted to be under their control^23^. Interestingly, super-enhancer domains seem to anchor regions lying on different chromosomes, thus distinguishing such higher order structures from the loop extrusion structures organized by cohesin which create only intrachromosomal links^23^. This is in line with our analysis performed for inter-chromosomal Hi-C contacts, which showed that loci overlapping super enhancers engage in more Hi-C contacts (**Figure 4E**). Same analysis also showed that HIV-1 targeted loci with super-enhancers more frequently engage in inter-chromosomal contacts with other HIV-1 loci not overlapping super enhancers, when compared to loci without super enhancers or without HIV-1 integration.This could explain part of HIV-1 targets that do not linearly overlap with super enhancers. A question that remains open is whether insertion into clusters of super-enhanced domains is mediated by the HIV-1 integrase and whether it confers a selective advantage to the virus. Among the super-enhancer proteins that could play a role in directing the viral genome towards the super-enhanced regions of the cellular genome, P300 and BRD4 seem to be the most promising candidates. P300, a 300 kDa histone acetyltransferase (HAT) used to identify traditional^47, 48^ as well as super-enhancers^12, 49, 50^ is an interaction partner of the HIV-1 integrase. The catalytic activity of this HAT, acetylating different residues in the C terminal domain of the integrase was shown to be important for the DNA binding activity of the viral enzyme^51^. A plausible additional function of this interaction could be to target the viral genome to the super-enhanced regions of the cellular chromatin.

BRD4, member of the Bromodomain and extraterminal (BET) family of proteins, as one of the *bona fide* super-enhancer proteins, does not bind the HIV-1 integrase^52, 53^ but has a well established role in HIV-1 latency^54, 55^. The mechanism of action has recently been ascribed to the short isoform of BRD4, which recruits a repressive SWI/SNF complex to the viral LTR^56^. According to this study, unloading the short isoform, occurring rapidly upon JQ1 treatment, would leave the long isoform engaged in the transcriptional activation of the viral genome^56^. The same mechanism could account for the activation of cellular genes upon JQ1 treatment^11^. In fact, our RNA-Seq data show that genes with super-enhancers are both up- and down-regulated upon JQ1 (**Supplementary Figure 2B**). At the same time, genes targeted by HIV-1 are more responsive to JQ1 than non-HIV-1 targets (**Supplementary Figure 2C**). This implies that HIV-1 preferentially targets the genes that have a rapid and tightly regulated transcriptional response. Given the opposing role of BRD4 on viral LTR and cellular genes, insertion into the genes delineated with super-enhancers, might represent a source of transcriptional fluctuations^57, 58^.

In endothelial tissue, during proinflammatory stimulation, rapid clustering of inflammatory super-enhancer domains depend on p65 of NF-κB, which was shown to drive a global redistribution of BRD4^59^. Localization of BRD4 to chromatin was proposed to facilitate the transition between cellular states^59^. Our experiment on resting T cells, where JQ1 pretreatment was used prior to CD4^+^ activation with CD3/CD28 antibody suggest that postponed repositioning of genes towards the nuclear periphery might be a consequence of redistribution or unloading of BRD4 from chromatin (**Supplementary Figure 6F**). That this might be correlated to a rapid and global redistribution of p65 of NF-κB is an intriguing possibility that merits further examination. Along the same line, it has to be taken into account that PKC and NF-κB inhibitor, Bryostatin-1^60–62^ efficiently reactivates latent HIV-1. The same is true for JQ1, which reverts the latent viral state in cell lines and primary T cell models^54, 56^, as well as in samples from HIV-1 infected individuals^63, 64^. Interestingly, when used in combination these two drugs seem to have a particularly potent effect, further reinforcing the notion that HIV-1 integrates and persists in super-enhanced regions of the genome. While it can be argued that HIV-1 integration patterns into super-enhanced regions represent accidental events of selection of the first open chromatin regions at the sites of nuclear entry, one should also bear in mind that these sites of integration are those found in patients on cART. Hence, insertion into the genes at the nuclear periphery with here described specific properties might be evolutionary advantageous for viral persistence under antiretroviral treatment.

---

**Supplementary Figure 1**.

**A)** HIV-1 integration sites (IS) inside genes. Box plot represents percentage of IS inside genes for cART treated patients (n lists=6) in violet and in vitro infections (n lists=2) in blue. Red line depicts median while whiskers stretch from 5^th^ to 95^th^ percentile.

**B)** Analysis of number of unique genes containing integration sites with number of observed integrations. All integration data sets are sorted by decreasing size and cumulative number of integration sites is plotted on X-axis while Y-axis shows number of unique genes that have integrations. Number of unique genes found when analyzing different number of data sets linearly depends (adjR^2^ = 0.9849, p=7.72e^-08^) on the number of integrations in observed data sets.

**C)** HIV-1 recurrently integrates into a subset of genes. Bar plot represents number of genes (RIGs) shared among at least x different data sets. Number of RIGs shared among different data sets decreases exponentially as more data sets are taken into consideration.

**D)** Bar plot shows percentage of genes that have super-enhancer in proximity in group of genes without HIV-1 integrations (0 lists), on 1 list and in group of RIGs (2 or more lists).

**E)** ROC analysis represented in heatmap summarizing the co-occurrence density of integration sites and epigenetic modification obtained by ChIP-Seq for H3K27ac, H3K4me1, BRD4, MED1, H3K36me3, H4K20me1, H3K4me3, H3K27me3 and H3K9me2. HTLV, HIV-1 and MLV integration data sets are shown in the columns, and epigenetic modifications are shown in rows. Associations are quantified using the ROC area method; values of ROC areas are shown in the color key at the right.

**Supplementary Figure 2**.

**A)** mRNA expression profiles and protein levels of MYC upon JQ1 treatment (500nM JQ1 for 6 h). mRNA levels are normalized over GAPDH and mean and SD are derived from three independent experiments. Representative protein levels of c-Myc and actin are shown on Western blot.

**B)** Volcano plot change in mRNA levels for genes upon JQ1 treatment with respect to the vicinity to super-enhancers.

**C)** Bar plot shows percentage of genes that are downregulated, unchanged, and upregulated upon JQ1 treatment. Genes are grouped by number of lists they occur in and by presence or absence of super enhancer in either gene body or 5 kb upstream of a gene.

**Supplementary Figure 3.**

**A)** Comparison of CD4^+^ IS data sets with Jurkat B-HIVE IS data set. Fraction of genes from a data set that is shared with at least one other data set is shown on Y-axis. Bar plot shows that different data sets share most of targeted genes among each other, while randomly chosen subsets of genes (minRND and maxRND) are only partially shared with genes from other data sets.

**Supplementary Figure 4.**

**A)** Density plot of inter-chromosomal pairwise contact scores (see Methods) between super enhancers and HIV hotspots. HIV hotspots were identified within genomic bins of 100kbp, then sorted by count of HIV insertions and split in percentile groups, e.g. top 0.5% in HIV count, from 0.5% to 1%, etc. Super-enhancers show strongest inter-chromosomal contacts with other super-enhancers (observe the higher density at higher contact scores, red line), followed by contacts between super-enhancers and the most HIV-dense hotspots (pink line). The scores of interaction show a monotonous decay as the HIV hotspots are more sparse.

**Supplementary FIgure 5.**

**A)** Proportion of 3D sub-compartments covered by Jurkat chromatin features available from the literature. Coverage was computed as the span of enriched ChIP-Seq signal divided by the sub-compartment size.

**B)** Scatter plot of HIV density in gene bodies versus endogenous expression in Jurkat cells. Each dot represents a protein-coding gene. Dot colors identify the 5 different sub-compartments. Sub-compartments A1 and A2 show similar distributions of gene expression and almost identical effects of gene expression on HIV density (see slopes of linear models). However, A1 shows higher HIV density overall compared to A2 (see vertical shift of approximation curves), suggesting an intrinsic preference of HIV for A1 independent of the gene expression level.

**Supplementary Figure 6**.

**A)** Adjusted p value for change in expression of genes upon activation of CD4^+^ T cells. Genes are grouped by number of HIV-1 lists they appear in. Red dashed line represents adjusted p value of 0.05.

3D immuno-DNA FISH in resting and activated CD4^+^ T cells. Representative images of **B)** *PTPRD* and *TAP1*; **C)** *MYC*; **D)** *BACH2*, *NPLOC4* and *RPTOR*. **E)** Table summarizing spatial relationships on chromosome 11 and 17 in resting and activated CD4^+^ T cells. **F)** Effect of SE dismantling by JQ1 on the T cell activation-induced movement of RIGso: 3D immuno-DNA FISH images of *STAT5B* (left) and *GRB2* (right) in CD4^+^ T cells treated with 500 nM JQ1 or DMSO, and activated for 20 h: Green: gene, red: lamin B1, blue: DNA counterstaining with Hoechst 33342. Scale bars represent 2 μm. Cumulative frequency plots in the lower panels show combined data from both experiments (n = 100, black: resting cells, red: activated cells). P values obtained by KS tests are indicated.

## METHODS

### Primary cell isolation, culture, treatments and infection

For CD4^+^ T cells isolation, whole blood was mixed with RosetteSep Human CD4^+^ T cell enrichment cocktail beads according to the manufacturer instructions and CD4^+^ T cells were separated using Histopaque Ficoll gradient by centrifugation. Cells were cultured in complete T cell medium (RPMI-1640 + 10% FBS + primocin), left in resting state or activated with Dynabeads Human T-Activator CD3/CD28 and plated in complete medium supplemented with 5ng/ml IL-2 for 20 to 72 h at 37°C. Cells were treated when indicated with 500 nM JQ1(+) or DMSO for 6 hrs at 37°C. 1×10^6^ activated CD4^+^ T cells were infected with 0.5-1 µg of p24 of virus by spinoculation for 90 min at 2300 rpm at RT in the presence of polybrene at 37°C. Virus stocks were produced from the viral clone HIV-1NL4_3 and a mutant that harbors a frameshift (FS) mutation in the *env* gene (pNL4_3-envFS) and was pseudotyped with vesicular stomatitis virus glycoprotein (VSV-G), resulting in a frameshift virus that performs a single-round infection (HIV-1NL4_3 FS). Cells were then incubated for 72 h at 37°C. When indicated, 14 h after infection with HIV-1NL4_3, cells were treated with the fusion inhibitor T20 to prevent multiple infection and integration. All viral stocks were generated by transfecting viral DNA in HEK 293T cells and collecting supernatants after 48 to 72 hours following sucrose gradient purification of virus articles. Viral production was quantified in the supernatants for HIV-1 p24 antigen content using Innotest HIV antigen mAB kit (INNOGENETICS N.V. Gent, Belgium). The human Jurkat T cell line (obtained from the cell collection of the Center for Genomic Regulation, Barcelona) was grown at 37°C under a 95% air and 5% CO2 atmosphere, in RPMI 1640 medium (Gibco) supplemented with 10% FBS (Gibco), 1% penicillin-streptomycin (Gibco) and 1% GlutaMAX (100x) (Gibco). Jurkat cells were passaged every 2 d with a 1:5 dilution. Cells were tested for mycoplasma yearly.

### Linear amplification-mediated PCR (LAM-PCR) to map HIV-1 insertion sites in primary cells

The mapping was performed as described previously^65, 66^ using 1µg of genomic DNA from HIV-1 NL4-3 infected primary human CD4^+^ T cells.

### Inverse PCR to map HIV-1 insertion sites in primary cells

The mapping of HIV was performed based on the protocol published by Chen *et al.*^67^ with modifications. Briefly, 3 µg genomic DNA from HIV-1NL4_3-infected CD4^+^ T cells treated with 500 nM JQ1 or DMSO before infection were digested by 2 µL 10,000 U/mL AluI (NEB, R0137S) and 2 µL 10,000 U/mL BglII (NEB, R0144S) in NEBuffer 2.1 in 50 µL final volume at 37 ℃ for 3 hours. The reaction was heat-inactivated at 80 ℃ for 20 min. BglII digestion aims to eliminate byproducts, which contain only the sequence of the HIV-1 backbone after AluI digestion. The double-digested products were diluted in 1 mL T4 DNA ligase buffer, then self-ligated by adding 2 µL 30 U/µL T4 DNA ligase (Thermo Fisher Scientific, EL0013) and incubating at 16 ℃ overnight. The ligation reaction was ethanol-precipitated the following day. The pellet was resuspended in 84 µL distilled water. To destroy non-circularized genomic DNA, 4 µL 25 mM ATP and 2 µL 10 U/µL Plasmid-Safe™ ATP-Dependent DNase (Epicentre, E3101K) were added with 10X Reaction Buffer in 100 µL final volume at 37 ℃ for 2 hours. The reaction was heat-inactivated at 70 ℃ for 30 min.

6 µL Plasmid-Safe-digested products were mixed in 50 µL standard Phusion polymerase reaction mix (Thermo Fisher Scientific, F530S) in GC buffer, with 0.1 µM primers GAT1786 (annealing to the Illumina PE1.0 primer) and one indexing primer GAT-int_5LTR (annealing to the 5’ end of the LTR) or 0.1 µM primers GAT1799 (annealing to the Illumina PE1.0 primer) and one indexing primer GAT-int_3LTR (annealing to the 3’ end of the LTR) for each condition of the sample. The cycling conditions were as follows: 98 ℃ for 1 min; 98 ℃ for 20 sec, 55 ℃ for 1 min, 72 ℃ for 5 min (2 cycles); 98 ℃ for 20 sec, 62 ℃ for 1 min, 72 ℃ for 5 min (27 cycles); 72 ℃ for 5 min. GAT-int_5LTR and GAT-int_3LTR primers add the Illumina PE2.0 primer and a 6-nucleotide index to the amplicons. PCR products ran as a smear on agarose gel. The primers used are described in **Additional file 3**.

### Fluorescence *in situ* hybridization (FISH)

Approximately 3×10^5^ CD4^+^ T cells were plated on the PEI coated coverslips placed into a 24-well plate for 1 hour at 37 C. Cells were treated with 0.3x PBS to induce a hypotonic shock and fixed in 4% PFA/PBS for 10 min. Coverslips were extensively washed with PBS and cells were permeabilized in 0.5% triton X-100/PBS for 10 min. After three additional washings with PBS-T (0.1% tween-20), coverslips were blocked with 4% BSA/PBS for 45 min at RT and primary antibody anti-lamin B1 (1:500 in 1% BSA/PBS) was incubated overnight at 4°C. Following three washings with PBS-T, fluorophore-coupled secondary antibody (anti-rabbit, coupled to Alexa 488, Alexa 568 or Alexa 647, diluted 1:1000 in 1% BSA/PBS) were incubated for 1 h at RT, extensively washed and post fixed with EGS in PBS. Coverslips were washed three times with PBS-T and incubated in 0.5% triton X-100/0.5% saponin/PBS for 10 min. After three washings with PBS-T, coverslips were treated with 0.1 M HCl for 10 min, washed three times with PBS-T and additionally permeabilized step in 0.5% triton X-100/0.5% saponin/PBS for 10 min. After extensive PBS-T washings, coverslips were equilibrated for 5 min in 2x SSC and then put in hybridization solution overnight at 4°C. For the HIV-1 FISH, RNA digestion was additionally performed beforehand using RNAseA (100ug/ml).

#### FISH without IF for HTI

1-2×10^6^CD4^+^ T cells in 500 ul of medium were adhered to coverslips by centrifugation at 350 g for 10 min at RT. The coverslips were washed in PBS and cells were fixed in 4% PFA/PBS for 10 min followed by extensive PBS washing. Permeabilization was performed by incubation in 0.5% triton X-100/0.5% saponin/PBS for 20 min. After three washings with PBS, cells were treated with 0.1 M HCl for 15 min. Coverslips were washed twice for 10 min with 2x SSC, put in hybridization solution overnight at 4°C.

#### DNA probe labeling

BAC or PAC DNA was extracted using a Nucleobond Xtra Maxiprep kit according to the manufacturer’s instructions, while HIV-1 plasmid HXB2 was purified using Qiagen plasmid extraction kit. FISH probes were generated in a Nick translation reaction using three different protocols.

##### (1) Conventional BAC labeling with digoxigenin (DIG)-coupled dUTPs

3 μg of BAC DNA were diluted in H2O in a final volume of 16 μl. 4 μl of DIG-Nick translation mix (Roche) were added and the labeling reaction was carried out at 15°C for up to 15 h. The labeling reaction was performed by using a fluorophore-coupled dUTPs in the same concentration as biotin-16-dUTP in *(2*)

##### (2) Conventional HIV-1 labeling with biotin-coupled dUTPs

For HIV-1 labeling, a biotin-dUTP nucleotide mix containing 0.25 mM dATP, 0.25 mM dCTP, 0.25 mM dGTP, 0.17 mM dTTP and 0.08 mM biotin-16-dUTP in H2O was prepared. 3 μg of pHXB2 were diluted with H2O in a final volume of 12 μl, and 4 μl of each nucleotide mix and Nick translation mix (Roche) were added. Labeling was performed at 15°C for 3 to 6 h.

##### (3) Direct labeling with fluorescently labeled dUTPs

For dual-color FISH or improvement of signal-to-noise ratio in single-color FISH, probes were labeled using the fluorophore-coupled nucleotides SpectrumGreen dUTP (Abbott), SpectrumOrange dUTP (Abbott) and Red 650 dUTP (Enzo).

#### Abbott Nick translation kit

1 to 3 μg of BAC DNA were diluted in a final volume of 22.5 μl H2O. 2.5 μl of 0.2 mM fluorophore-coupled dUTP, 5 μl of 0.1 mM dTTP, 10 μl of a dNTP mix containing 0.1 mM of each dATP, dCTP and dGTP, and 5 μl of 5x Nick translation buffer (Abbott) were added and reagents were mixed well by vortexing. The reaction was started by addition of 5 μl Nick translation enzymes (Abbott) and incubated at 15°C for 13 to 14 h.

After labeling reactions, the probes were checked for their size on a 1% agarose gel, and 200 to 500 bp probes were purified using Illustra Microspin G-25 columns according to the manufacturer’s instructions. Probes were precipitated with Cot-1 and herring sperm DNA, 3 M sodium acetate ice-cold 100% ethanol at −20°C overnight.

After 70% ethanol wash the pellet was air-dried for several hours, resuspended in formamide and incubated shaking at 37°C for 15 min to dissolve the pellet. Finally, probes were mixed with 10 μl of 4x SSC/20% dextran sulfate, denatured at 95°C for 5 min, kept on ice for 2 min and then stored at −20°C.

#### Probe hybridization and detection

1-6 µl of probe was loaded on glass coverslips and heat denatured in metal chamber at 80^0^C for 8min in a water bath. Hybridization was carried out for 48 hours at 37^0^C. 4 washings in 2XSSC (10 min each) at 37^0^C were followed with 2 washings in 0.5X SSC at 56^0^C.

FISH development for DIG-labeled BACs was performed by using FITC-labeled anti-Digoxigenin antibody (Roche), whereas biotin-labeled HIV-1 probes were detected by TSA Plus system from Perkin Elmer, that allows significant amplification of the signal, by using an anti biotin antibody (SA-HRP) and a secondary antibody with a fluorescent dye (usually FITC for HIV).

For the directly labeled probes after initial washings nuclei were stained with Hoechst 33342 (1:5000 in PBS), washed in PBS and then mounted using mowiol.

### Microscopy and image analysis

#### (1) Classical confocal microscopy and manual image analysis

Three-dimensional stacks were acquired with a Leica TCS SP8 confocal microscope using a 63x oil immersion objective. Distance measurements were performed using Volocity (Perkin Elmer). The smallest distance between the FISH signal and the nuclear lamina, stained by immunofluorescence for lamin B1, was determined, and measurements were normalized to the nuclear radius (defined as half of the maximum diameter of the lamin B1 ring). Signal-to-radius ratios were either binned into three classes of equal volume (zones 1 to 3)^6^, or plotted on a cumulative frequency plot. Kolmogorov-Smirnov (KS) tests were performed to compare the distributions of positioning of a gene between two conditions (resting vs. activated or DMSO vs. JQ1).

#### (2) High-throughput imaging (HTI) and image analysis of dual-color FISH

Images were acquired with a spinning disk Opera Phenix High Content Screening System (PerkinElmer), equipped with four laser lines (405 nm, 488 nm, 568 nm, 640 nm). Images of FISH experiments to calculate 3D distances were acquired in confocal mode using a 40X water objective lens (NA 1.1) and two 16 bit CMOS cameras (2160 by 2160 pixels), with camera pixel binning of 2 (corresponding to 299 nm pixel size). For each sample, 11 *z*-planes separated by 0.5 µm were obtained for a total number of at least 36 randomly sampled fields, which acquired per condition a minimum of 16×10^3^ cells. Image analysis was performed using the Harmony high-content imaging and analysis software (version 4.4, Perkin Elmer), using custom made image analysis building blocks. Nuclei were segmented based on the Hoechst nuclei staining signal of maximum projected images using the algorithm B and cells in the periphery of the image were excluded from further analysis. FISH probe detection was performed by using the spot detection algorithm C and custom made scripts were used to calculate the Euclidean distances between all the different colored probes per cell. Single cell level data were then exported and custom made R scripts were used to select the minimum distance between the different FISH probes per allele basis. To exclude spurious spot detection events from the analysis, only the distances of cells with two FISH probes detected per channel were calculated and plotted (Graph Pad, Prism).

### Quantitative real-time PCR (qPCR)

Up to 5×10^6^ CD4^+^ T cells were used for RNA extraction with InviTrap Spin Kit (Stratec Biomedical) according to the manufacturer’s instructions and up to 500 ng of RNA was retro-transcribed using Moloney murine leukemia virus reverse transcriptase (M-MLV RT) from Invitrogen according to the manufacturer’s instructions. Gene expression analysis were performed in duplicates using IQ supermix from Biorad in CFX96/C1000 Touch Real-Time PCR system, as described in Lusic et al., 2013. Statistical analysis of qPCR data was performed using Graphpad. Taqman assays used: for MYC Taqman Hs00153408_m1 FAM/MGB, and for GAPDH 4310884E VIC/TAMRA.

### Western blotting

5×10^6^ cells were harvested and homogenized in lysis buffer (20 mM Tris-HCl, pH 7.4, 1 mM EDTA, 150 mM NaCl, 0.5% Nonidet P-40, 0.1% SDS, 0.5% sodium deoxycholate supplemented with protease inhibitors (Roche) for 10 min at 4°C and sonicated (Bioruptor) for 5 min. Equal amounts of total cellular proteins (20 μg), as measured with Bradford reagent (Biorad), were resolved by 10% SDS-PAGE, transferred onto nitrocellulose membrane (GE Healthcare) and then probed with primary antibody, followed by secondary antibody conjugated with horseradish peroxidase. The immuno-complexes were visualized with enhanced chemiluminescence kits (GE Healthcare). Antibodies used: for MYC 9E10 SC-40 from Santa Cruz and for actin Anti-β-Actin AC-74 from Sigma Aldrich.

### Chromatin immunoprecipitation (ChIP)

20×10^6^ CD4+ T cells were washed 1 time in PBS prior to crosslinking with 1% Formaldehyde for 10 minutes at room temperature, followed by termination of the reaction with 125mM glycine on ice. Cell pellet was washed 2 times with PBS at 4C and was lysed in 0.5% NP-40 buffer (10mM Tris-Cl pH7.4, 10 mM NaCl, 3 mM MgCl_2_, 1 mM PMSF and Protease Inhibitors). For histone ChIPs, obtained nuclei were washed once in the same buffer without NP-40. Nuclei were resuspended in 0.5% NP-40 buffer supplemented with 0.15 % SDS and 1.5 mM CaCl2. Nuclei were incubated at 37 C for 10 minutes prior to addition of Micrococcal Nuclease (16 units of the enzyme), and the reaction was stopped after 7 minutes with 3 mM EGTA. DNA was additionally sheared by sonication (Covaris or Bioruptor, Diagenode) to an average size of DNA fragments below 500 bps. For BRD4 and MED1 ChIPs, nuclei were lysed in 1% SDS, 50 mM Tris pH 8 and 10mM EDTA and sonicated (Bioruptor, Diagenode) to an average size of DNA fragments below 500 bps. Extracts were then diluted up to 0.01% SDS, 1% Triton-X, 20mM Tris pH 8, 150mM NaCl 2 mM EDTA. Extracts were precleared by 1 hr incubation with protein A/G Magna ChIP beads at 4C and diluted with 5x IP buffer to a final concentration of 140 mM NaCl and 1 % NP-40. Lysate corresponding to 3-4 x 10^6^ million of cells was then incubated with 2-4 µg of the indicated antibody overnight at 4^0^ C, followed by a 2.5 hr incubation with Magna ChIP Protein A/G Magnetic Beads (Millipore). Beads were then washed thoroughly with RIPA150, with LiCl – containing buffer and with TE buffer, RNAse treated for 1 hr at 37^0^C, and Proteinase K treated for 2 hours at 56^0^C. Decrosslinking of protein–DNA complexes was performed by an overnight incubation at 65^0^C. Additional 1 hr of Proteinase K digestion was performed at 56^0^C and DNA was then extracted using Agencourt AMPure XP beads (Beckman Coulter) and quantified by real time PCR. The following antibodies were used ChIP: H3K27ac (ab4729), H3K4me3 (ab8580) H3K36me3 (ab9050), BRD4 (Bethyl lab A301-985A100), MED1 (Bethyl lab A300-793A), IgG Rabbit (ab46540).

### ChIP-Seq and RNA-Seq

ChIP-Seq: Approximately 10 ng of corresponding inputs and ChIP-ed DNA from primary CD4 T cells: H3K27Ac, H3K4me3, H3K36me3, H4K20me1 and H3K9me2 Ips was prepared for the sequencing using NEBNext^®^ Ultra™ II DNA Library Prep Kit for Illumina^®^ RNA-Seq: 5×10^6^ DMSO and 500 nM JQ1 treated CD4^+^ T cells from three independent donors were used for RNA extraction with InviTrap Spin Kit (Stratec Biomedical) according to the manufacturer’s instructions and libraries for sequencing were prepared by using rRNA depletion kit NEBNext^®^ and NEBNext^®^ Ultra™ RNA Library Prep Kit for Illumina^®^. Sequencing was performed with 2 × 75 bp read length on the NextSeq platform.

### In situ Hi-C protocol

Hi-C was performed based on the protocol published by Rao *et al*.^68^ with modifications. Briefly, one million cells were crosslinked with 1% formaldehyde for 10 min at room temperature with gentle rotation. Nuclei were permeabilized by 0.25 mL freshly prepared ice-cold Hi-C lysis buffer [10 mM Tris-HCl pH8.0, 10 mM NaCl, 0.2% Igepal CA630 (Sigma, I8896-50ML) and 1x Roche complete protease inhibitors (Roche, 11836153001)]. DNA was digested with 100 units of MboI (NEB, R0147M) at 37 °C overnight, and the ends of digested fragments were filled in by using 0.4 mM biotinylated deoxyadenosine triphosphate (biotin-14-dATP; Life Technologies, #65001) and ligated in 1 mL by incubating at 24 °C overnight with gentle rotation. After reversal of the crosslinks, ligated DNA was purified and sheared to a length of 400 bp. Ligation junctions were pulled down with 75 μL of 10 mg/mL streptavidin C1 beads. 10 μL of DNA-on-beads were amplified in 50 μL standard Herculase II Fusion DNA Polymerase reaction mix (Agilent Technologies, #600675) with 1 μM NEBNext Universal primer and index primer (NEB, E6040S). The cycling conditions were as follows: 98 °C for 2 min; 98 °C for 20 s, 65 °C for 30 s and 72 °C for 45 s (8 cycles); and 72 °C for 3 min. PCR products were purified with 1.0x Agencourt AMPure XP beads (BECKMAN COULTER, A63880). Libraries ran as a smear on 1.5% agarose gel and the estimate the size of a smear was around 300 bp. The quality of the Libraries was assessed by digesting with ClaI (NEB, R0197S) and checking that the smear shifts downwards.

### Bioinformatic analysis

#### Integration sites

We analyzed eight lists of HIV-1 integration sites, five of them (^18, 19, 31–34^ were downloaded from retroviral integration database (RID)^42^. One was downloaded directly from^30^ and one was provided by Lusic lab (previously unpublished). We used only unique integrations from each study. If location of the integration was not precisely defined (spanning more than one nucleotide), we used midpoint as the location for that integration. All sites were converted to hg19 (GRCh37) version of the genome using R^69^ rtracklayer package,^69^. Gene coordinates were downloaded from Ensembl, GRCh37, February 2014. UCSC symbols were used for genes in UCSC (hg19), while others were named after ENSG identifier. We counted overlaps between integration sites and all GRCh37 genes disregarding strand and orientation. Gene coordinates and number of HIV integration lists can be found in **Additional file 2.**

#### Redefinition of RIGs

To each gene, we added a number representing number of lists that found HIV-1 integration inside this gene. We define recurrent integration genes (RIGs) as genes for which we found HIV integration in more than 2 data sets. To assess the relationship between number of integration sites included in analysis and number of genes discovered to have integrations, we did the following: We made lists of genes found to have integrations in each study. Next, we sorted those lists in decreasing order, by number of integration sites found in a study. We plotted cumulative sum of number of integrations found in studies on x-axis, and number of genes targeted in that study and not in any other studies before that study on y axis. We used linear regression to model this relationship.

#### ChIP-Seq data analysis

We analysed ChIP-Seq datasets obtained from this study (H3K4me3, H3K36me3, H3K27Ac, H4K20me1 and H3K9me2) and publicly available data sets from University of Washington Human Reference Epigenome Mapping Project H3K4me1 (SRA accession number: SRX342315), H3K27me3 (SRX342313), input for CD4 primary T cells (SRX252742). Datasets for BRD4 (SRR2971477), MED1 (SRR2971478) and corresponding Input (GSM1527712) were downloaded from^70^.ChIP seq reads were mapped to human genome (GRCh37) using Bbmap(^71^) with parameters minid=0.98, qtrim=lr, minavgquality=20. Resulting bam files belonging to same experiments were merged and sorted using bamtools. Average binding profiles in reads per million across sets of genes were made using ngsplot^72^. Peaks were called using MACS2^73^, for every dataset versus its matching input, with parameters --broad --broad-cutoff 0.1 -p 1e-9 -g 2.7e9 -B. All results were transformed to RPKM for downstream analysis and visualization.

For Jurkat cells, ChIP-Seq reads were mapped to hg19 using BWA-mem. BWA options were as follows: ‘-k17 -r1.3 -B2 -O4 -T22’ for read lengths less or equal to 30 nt, ‘-k18 -B3 -O5 -T28’ for read lengths less or equal to 40 nt and default options for longer reads. ChIP-Seq enriched regions were discretized using Zerone^74^ with mapping quality cutoff 20 and enrichment confidence 0.99. We used publicly-available ChIP-Seq profiles for Jurkat cell line. All datasets were obtained from NCBI Gene Expression Omnibus with the following series accessions: ERG and GABPA (GSE49091)^75^; H3K27Ac, CDK7 and PolII (GSE50622, GSE60027)^76^; H3K36me3, H3K79me3, H3K4me1, H3K9me3, H3K27Ac, H3K4me3, PolII, S5P and S2P (GSE65687)^77^; NRSF (GSE53366)^78^; PolII and CDK12 (GSE72023)^79^; ETS1, CBP and RUNX (GSE17954)^80^; H3K27Ac (GSE51522)^81^; PolIII (GSE20309)^82^; H3K27Ac and H3K27me3 (GSE59257)^83^; H3K4me3, H3K27me3, H3K79me2 and PolII (GSE23080); PHF6 (GSE45864); KDM2B (GSE70624)^84^; RUNX1, GATA3, TAL1, LMO1, TCF3 and TCF12 (GSE29181)^85^; H3K27Ac, MED1 and MYB (GSE59657)^86^; H3K4me3 (GSE35583)^87^; Lamin (DamID, GSE94971)^88^; RUNX1 (GSE42575)^89^; TAL1 (GSE25000)^90^; H3K4me3 and H3K79me2 (GSE60104)^91^; PolII (GSE25494)^92^; RUNX1, GATA3, H3K27Ac and CTCF (GSE68976)^93^; MYC, BRD4 and CDK7 (GSE83777); RUNX1 and GATA3 (GSE76181)^94^; CTCF (GSE12889)^95^; H2AX (GSE25577)^96^; UTX (GSE72300)^90^; YY1 (GSE99521)^97^. The list of ChIP-Seq data used in this study are in **Additional file 4.**

#### Super-enhancer calling

We used super-enhancer data for all activated CD4^+^ cell types CD4p_CD25-_Il17p_PMAstim_Th17 and CD4p_CD25-_Il17-_PMAstim_Th from dbSuper^41^. To define super-enhancers using our own ChIP-Seq data we followed the same procedures as in dbSuper. Thus, we defined super-enhancers using HOMER software^40, 41^ findPeaks with default parameters (’-style super’) on our H3K27Ac peaks (peak finding described above). Briefly, peaks found within a distance of 12.5 kb were stitched together into larger regions. Super-enhancer signal of each region was determined by the total normalized number of reads subtracted by normalized number of reads in the input peaks. Regions are sorted by score and super-enhancers are identified as regions with score higher than that defined by slope greater than 1. We defined a gene to be “proximal” to super-enhancer if it overlaps with one, or if we can find a super enhanced element 5 kb upstream of transcription start site.

#### Hi-C contacts

Hi-C reads were mapped using BWA-MEM with the following options: ‘-P -k17 -U0 -L0,0 -T25’. Each read end was mapped independently. Genuine Hi-C contacts were validated with the Hi.C pipeline (https://github.com/ezorita/hi.c), using the following discard filters: (i) contact pairs with mapping quality below 10, (ii) self-circularized molecules and (iii) reads with inferred insert size greater than 2000 bp after digestion and ligation. Hi-C contacts were then binned at 5kb resolution and stored in HDF5 format using Cooler (https://github.com/mirnylab/cooler).

#### Hotspots and HIV-dense genes

HIV genome-wide hotspots were identified dividing the genome in bins of 100 kbp and sorting the bins by HIV insertion count. HIV density in genes followed a similar rationale but the bins were designed to match gene bodies, as described by ENSEMBL GTF GRCh37 release 75. HIV density was computed as the number of insertions per kbp of gene body.

#### Identification of 3D sub-compartments

To effectively reveal the underlying patterns of the Hi-C matrix, the operations below were performed in order to normalize the matrix for a robust clustering process. First, observed-over-expected balancing was applied to remove off-diagonal effects^98^. Outliers, such as enhancer-promoter loops, were smoothed by thresholding the greatest values of the matrix to the 90-th percentile. On the balanced matrix, the correlation matrix was computed and its diagonal was set to 0. The outliers were further smoothed by applying a linear scaling from the 5-th and 95-th percentiles to −1 and +1, respectively. Spatial clusters were identified by k-means (with 10 restarts) to the first k=15 weighted eigenvectors, *i.e.* the 15 leading eigenvectors, each weighted by their respective eigenvalue. This process was repeated independently for each chromosome, delineating 15 spatial clusters per chromosome. Next, the normalized interchromosomal scores between two chromosomal sub-compartments were computed as the total number of Hi-C reads between them, divided by the product of their sizes. The five global sub-compartments were finally identified by k-means clustering of the k=5 leading weighted eigenvectors of the normalized interchromosomal scores. Compartment names A1, A2, AB, B1 and B2 were assigned based on their distribution on the AB score scale. A1 and A2 had strong and moderate enrichment in active transcription marks, respectively. AB, B1 and B2 showed very low levels of active marks and moderate to strong enrichment in H3K27me3, H3K9me3 and Lamin, respectively.

#### AB score

AB scores were derived from the first eigenvector of the Hi-C correlation matrix (as in Identification of 3D sub-compartments). We chose the reference A and B regions to be the most dissimilar 3D structures, i.e. the genomic bins with 10% top and bottom values of the first eigenvector, respectively. For each row of the correlation matrix we computed A_score_ and B_score_ as the sum of its values in the A and B reference regions, respectively. Finally, the AB score was computed as:

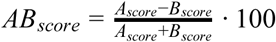

Yielding values between 100 for A-like regions and −100 for B-like regions. Ambiguous regions that are equally in contact with the reference A and B regions, or that are not in contact with them at all, will have AB scores close to 0.

### Pairwise contact score

Throughout the paper we used pairwise contact scores to quantify the amount of 3D interaction between many loci. Such interactions were compared pairwise and only between loci in different chromosomes. Considering only interchromosomal contacts we avoided the intrinsic bias produced by 1D short-range interactions. The pairwise score as the amount of Hi-C contacts within the interchromosomal region covered by the two loci divided by the product of their lengths in kbp. The units of this metric (#Hi-C reads/kb^2^) measure densities of contacts in a 2D region, allowing for fair comparison of different loci even if their spans are different.

### Statistical analysis

#### Generation of random matched control sites

We generated 10 control sites for every integration site in the following way: First we generated random 100 million numbers from 1 to largest chromosome length with seed set to 23779. Next, we generated 100 million chromosome names, where names were chosen at random but with weights corresponding to number of occurrences of each chromosome in our data set. This way we generated 100 million random possible positions for controls. Next, we excluded all the positions from this random set that were found in the blacklisted area of human GRCh37 genome^99^. Sites are available on request. We calculated distance to nearest gene for each possible integration site and divided them to subsets of 1000 base pair bins based on those distances. For each true integration site, we extracted a subset of all random possible positions that are located in the same bin of distance to their nearest gene as the integration site is to its nearest bin. Then, from those equidistant subset of random integration sites we randomly picked 10 to represent random matched controls for each real integration site.

#### Adding genomic feature value to integration sites and comparing integration sites to its random matched controls

First, we cut the genome into tiles of length 1000 base pairs and excluded blacklisted areas. We calculated 75^th^ quantile of RPKM values of each genomic feature (for each chromatin mark and transcription factor separately) over each tile. For super-enhancers we used 1 if super enhancer exists in a tile, and 0 if it does not instead of RPKM values. We assigned the value of the bin in which integration site is located to each integration site (and each random matched control). Next, we compared the value for each integration site only with its matched controls to determine the proportions of controls whose values equaled or exceeded that of the integration site. We scored each integration site in the following way: if this value was higher on true integration site than on matched control, we counted it as 1. If the value for the matched control was equal to true value on integration site, we counted it as ½, and if the value was lower, it was counted as 0. Final score for each integration site was calculated as average of those 10 values. Finally, we calculated empirical ROC area under the curve as average of all values for integrations in a data set. At last, we repeated this analysis on various bin sizes; 1Kb, 2Kb, 5Kb, 10Kb, 20Kb, 25Kb, 50Kb, 100Kb and on all genomic features and data sets and created a heatmap for every bin size (data not shown). We implemented p value calculation from^100^ in R. Briefly, all comparisons utilize the Wald-test statistic and are referred to a Chi Square distribution to obtain p-values.

### RNA-Seq data analysis

We mapped reads from RNA-Seq experiments to human genome (GRCh37 assembly, GENCODEV19) using BBMap with parameters maxindel=200000 xstag=unstranded ambiguous=random xmtag=t scoretag=t pairedonly=t minavgquality=20 maqb=51 minid=0.91. To calculate mean expression over replicates, we used rlog transformation from DESeq2 package^101^ to normalize the counts over all replicates, and calculated mean over all replicates. Differential expression for JQ1 treated cells VS control cells was done with the same package, following the Bioconductor RNA-seq workflow^102^.

For Jurkat cells, gene expression levels were derived from mRNA-seq experiments in Jurkat^103^. Sequencing reads were mapped to protein-coding Ensembl cDNA assembly GRCh37 release 75 using kallisto^104^ with options ‘–single’ (single-end mode), ‘–bias’ (sequence bias correction, ‘–s300’ (fragment length 300 nucleotides) and ‘–l100’ (s.d. 100 nucleotides). The counts of the different isoforms were summed to make a total count per gene copy in transcripts per million reads (tpm).

The list of RNA-Seq data used in this study are in **Additional file 5.**

## ACKNOWLEDGMENTS

We would like to thank the Infectious Diseases Imaging Platform (IDIP) and the platform coordinator Vibor Laketa (DZIF), as well as the genomics core facility of the CRG for their technical support.

This work was supported by German Center for Infection Research (DZIF) Thematic Translational Unit HIV-1 04.704 Infrastructural Measure to M.L, and to the Hector Grant “HiPNose: HiV Positioning in the Nuclear Space” to M.L and M.S.

We acknowledge the financial support of the Spanish Ministry of Economy and Competitiveness (‘Centro de Excelencia Severo Ochoa 2013-2017’, Plan Nacional BFU2012-37168), of the CERCA (Centres de Recerca de Catalunya) Programme / Generalitat de Catalunya, and of the European Research Council (Synergy Grant 609989).

