## Supplementary Materials for "Spatially clustered loci with multiple enhancers are frequent targets of HIV-1"

### Supplementary Figure 1

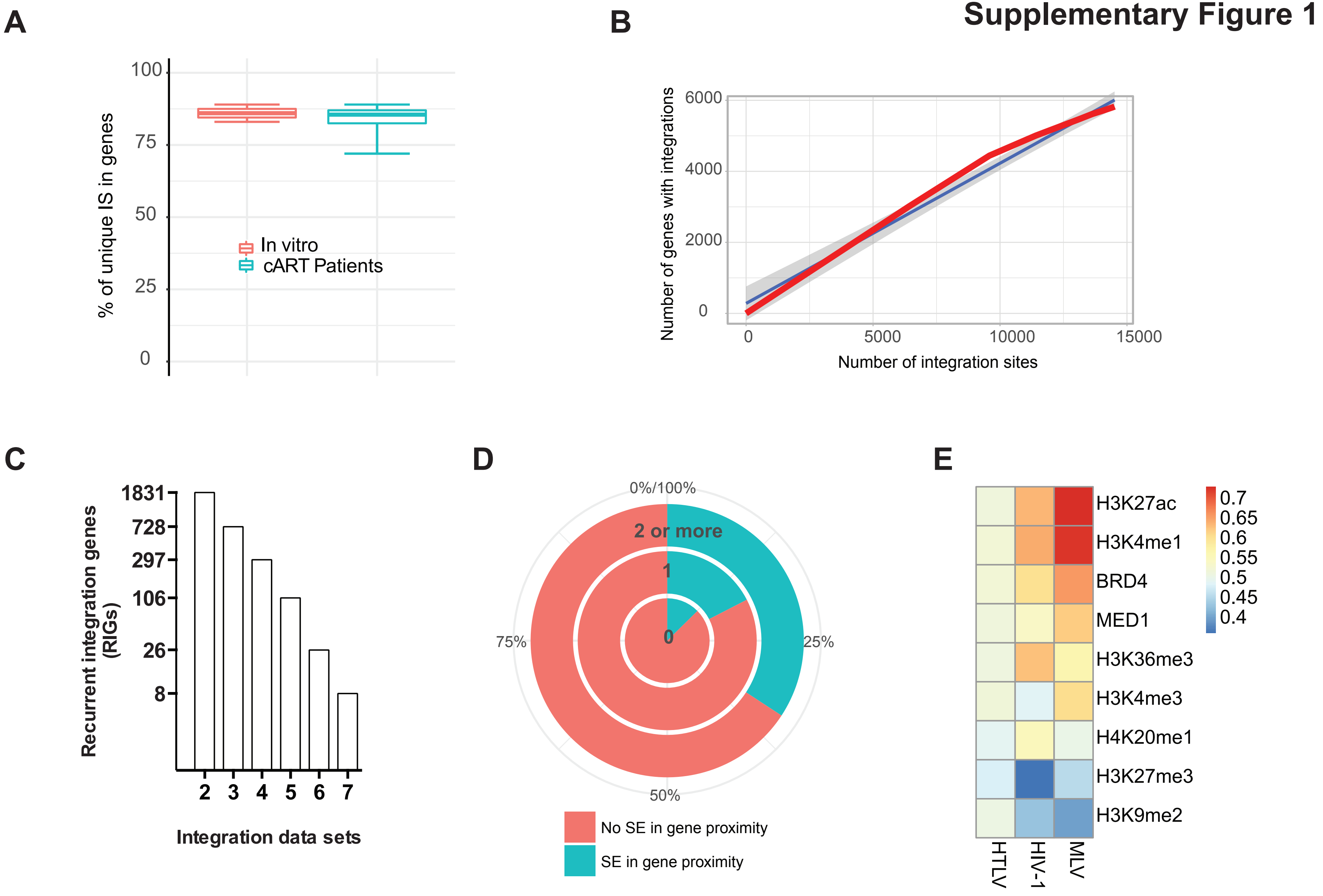

### Supplementary Figure 2

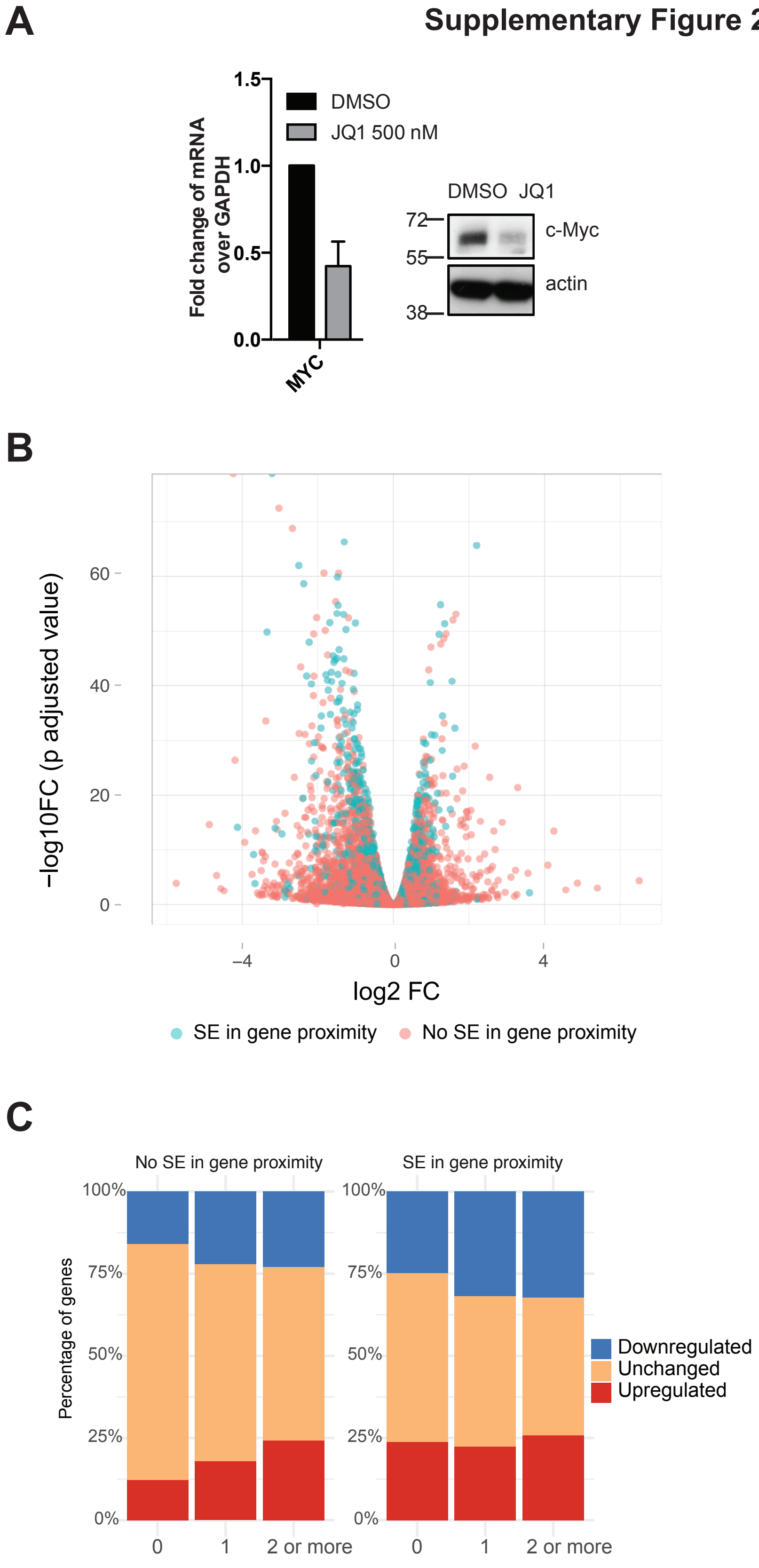

### Supplementary Figure 3

**A**

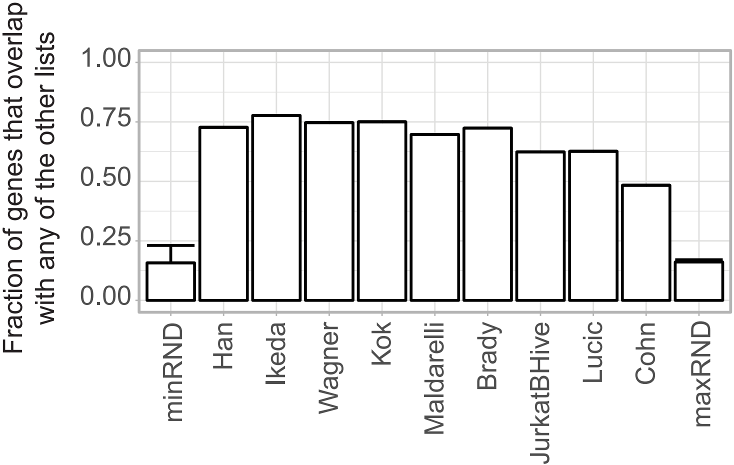

**Supplementary Figure 4**

**A**

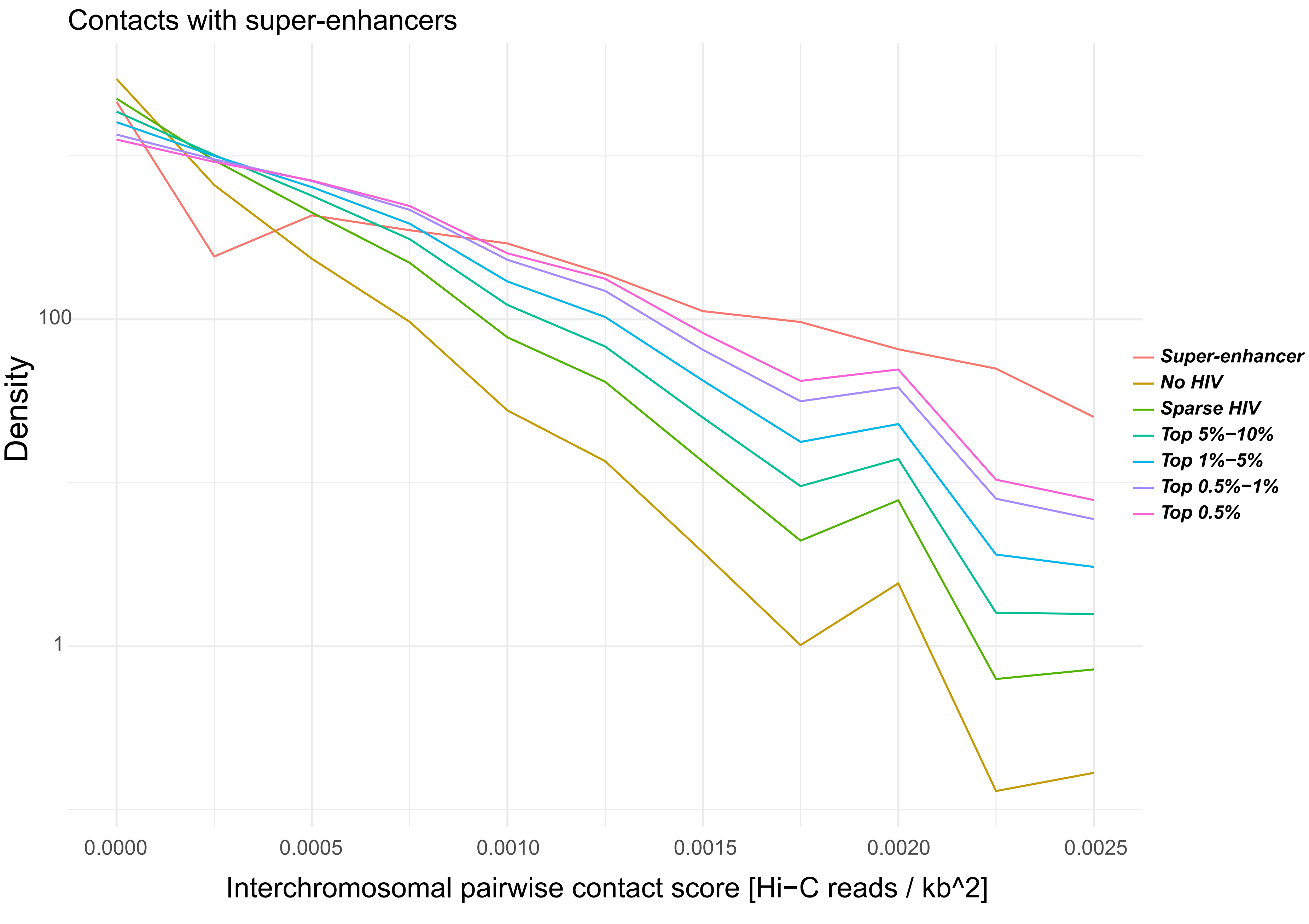

### Supplementary Figure 5

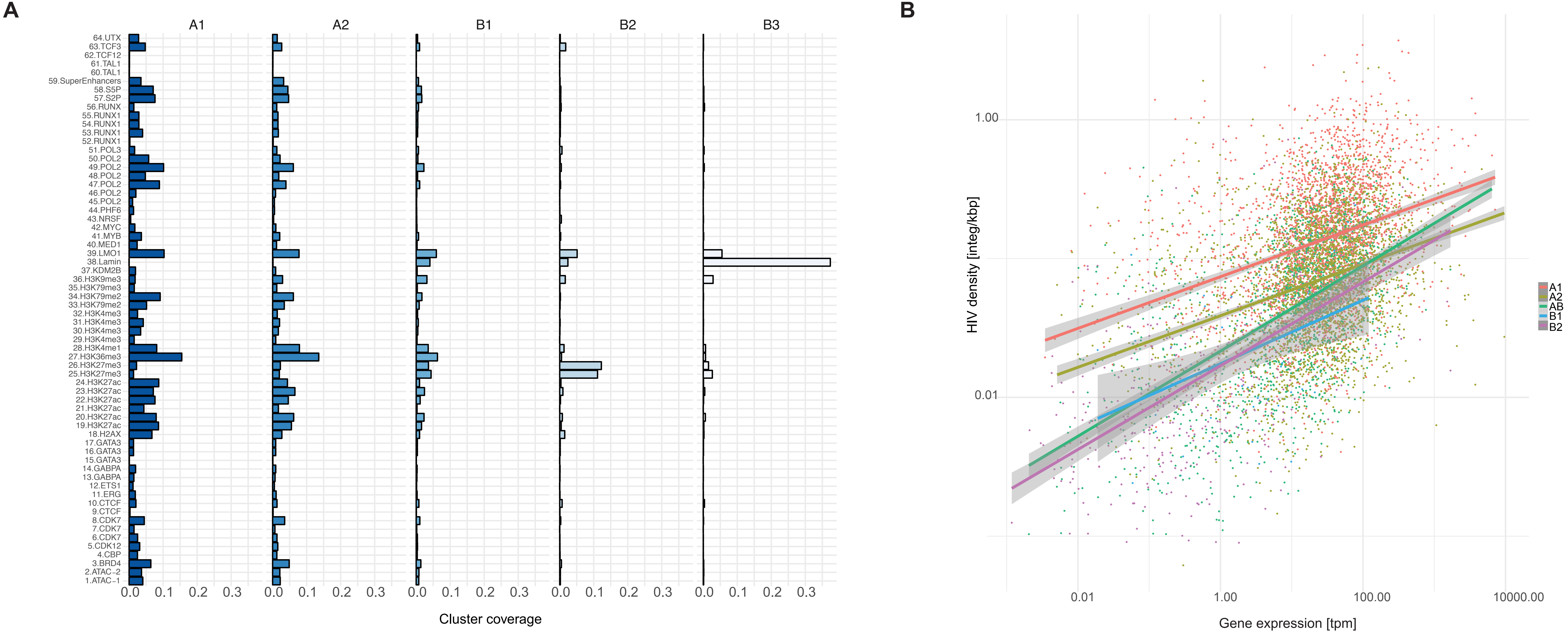

### Supplementary Figure 6

A

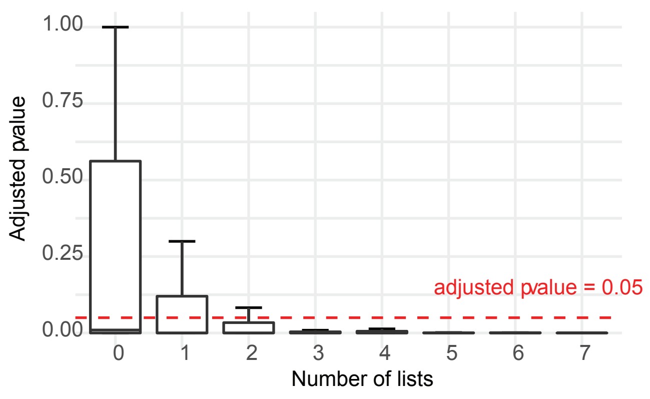


B

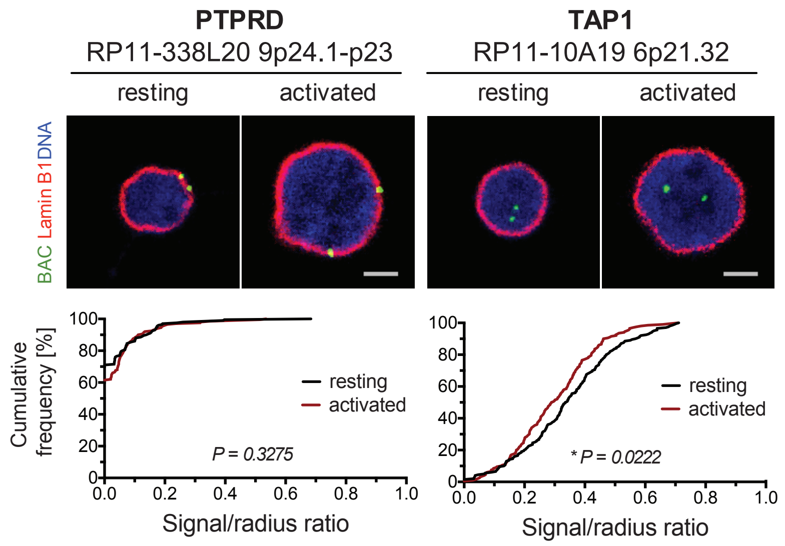


C

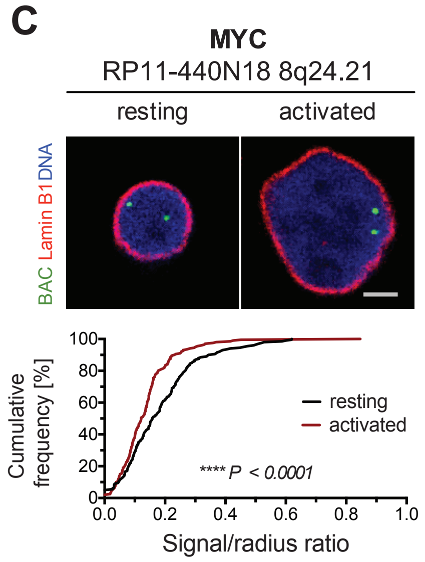


D

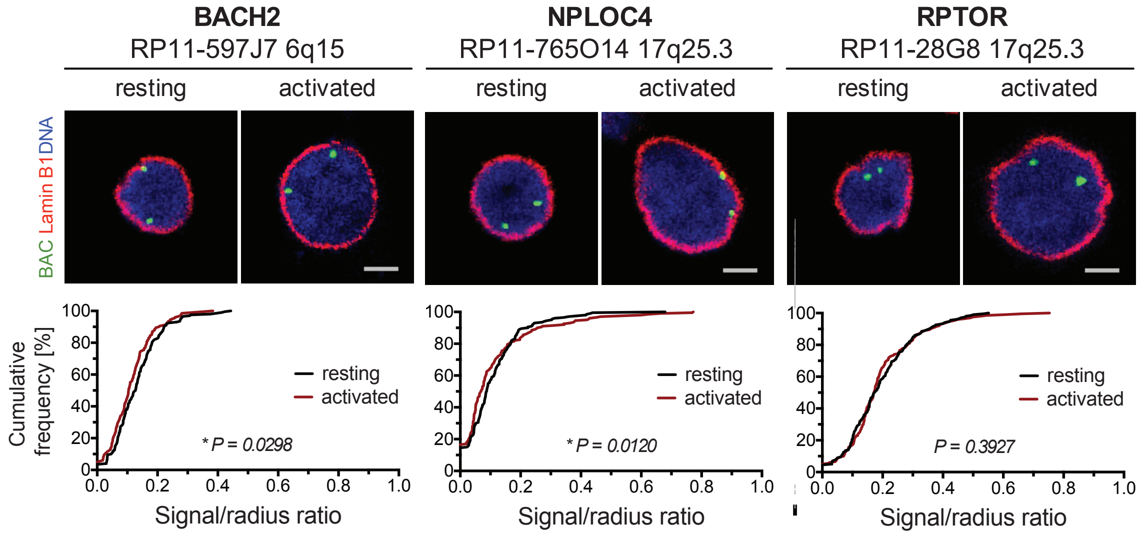


E

| RIGs Gene pair | Chromosome location | Distance (Mb) | Median distance (μm) | 75% percentile |
|---|---|---|---|---|
| PACS1 - KDM2A | 11q13.2 | 1.1 | 0.422849 | <0.668585 |
| | | | 0.422849 | <0.668585 |
| GRB2 - TNRC6C | 17q25.1 -3 | 2.6 | 0.598 | <0.945522 |
| | | | 0.598 | <0.945522 |
| RNF157 - GRB2 | 17q25.1 -3 | 0.8 | 0.422849 | <0.668585 |
| | | | 0.422849 | <0.598 |
| RPTOR - TNRC6C | 17q25.1 -3 | 2.7 | 1.495 | <2.11425 |
| | | | 1.495 | <2.15612 |
| GRB2 - RPTOR | 17q25.1 -3 | 5.3 | 1.61016 | <2.392 |
| | | | 1.495 | <2.15612 |
| NPLOC4 - RPTOR | 17q25.1 -3 | 0.8 | 1.81875 | <2.67434 |
| | | | 1.91453 | <2.75664 |

F

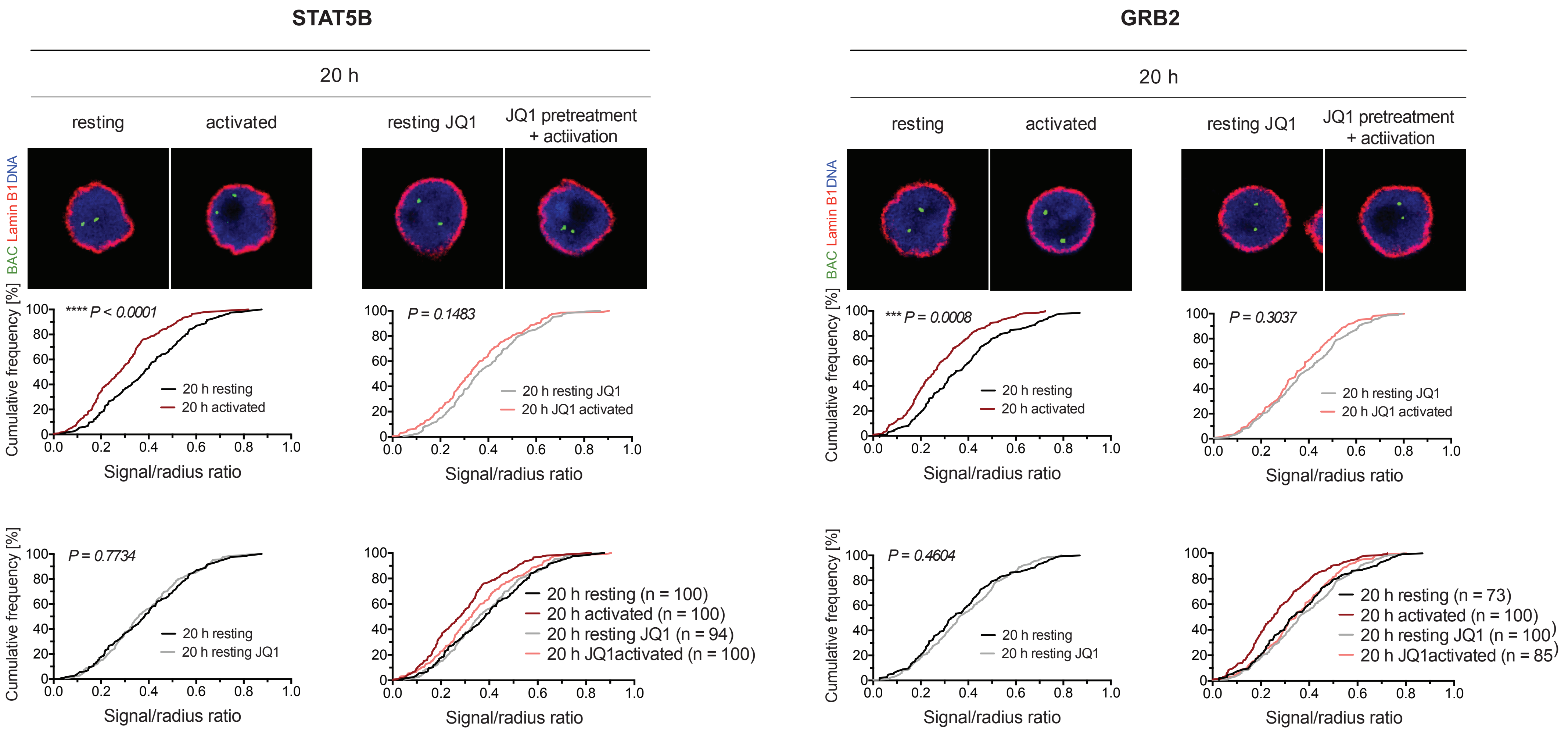
